# Control of Social Withdrawal of Mice Deficient for the Autism Gene *Magel2* by Restoration of Vasopressin-Oxytocin Dialogue in Septum

**DOI:** 10.1101/800425

**Authors:** Amélie M. Borie, Yann Dromard, Djodi Dufner, Emi Pollozi, Damien Huzard, Csaba Tömböli, Aleksandra Olma, Maurice Manning, Pascal Colson, Gilles Guillon, Françoise Muscatelli, Michel G. Desarménien, Freddy Jeanneteau

**Author notes:** Equal contribution (Michel G. Desarménien, Freddy Jeanneteau).

## Abstract

Intellectual and social disabilities are common comorbidities in adolescents and adults with *Magel2* gene deficiency characterizing the Prader-Willi and Schaaf-Yang neurodevelopmental syndromes. The cellular and molecular mechanisms underlying the risk for autism in these syndromes are unexplored. Here we used *Magel2* knockout mice combined with optogenetic/pharmacological tools to characterize disease modifications in the social brain network. We find that the degree of social novelty moderates a dialogue between vasopressin and oxytocin in the lateral septum, a region organizing sequential content of sensory experiences. Social withdrawal of mice lacking *Magel2* is alleviated by restoration of dialogue-lead by vasopressin. This preclinical study identifies the collective actions of vasopressin and oxytocin in the lateral septum as a key factor in the pathophysiology.

## Introduction

Autism Spectrum Disorders (ASD) affect 1 in 68 and are characterized by difficulties with communication, restrictive interests and repetitive behaviors influencing the ability to function properly (Diagnostic and Statistical Manual of Mental Disorders, 5th edition). Treatment of ASD is marginal such that management of disability-adjusted life years imposes substantial economic cost and burden on families and society^1^. Neurodevelopmental disorders characterized by mutations of chromosome 15q11-13 exhibit higher than normal risk for comorbid ASD, indicating the importance of genes in this locus for the pathophysiology^2,3^. It is a large chromosomal deletion of 15q11-13 in Prader-Willi Syndrome (PWS) and a disruption of one gene in this locus, *Magel2*, in Schaaf-Yang Syndrome (SYS) that provide respectively, 25% and 75% risk for ASD based on clinical assessment by an expert physician^2,3^. *Magel2* is a maternally imprinted, paternally expressed gene central to the pathophysiology of PWS and SYS^4^, which deficiency interferes with developmental functions essential for setting multi-scale organization of the nervous system controlling muscle tone and feeding^5,6^. Common symptomatic features of PWS and SYS are hypotonia, feeding difficulties during early life, social withdrawal, intellectual and/or developmental delay^4,7,8^.

Mice lacking *Magel2* gene are a good model of SYS and PWS with construct and face validity as knockout mice present hypotonia, feeding difficulties during early life and social deficits^9–11^. Treatment with oxytocin (OXT) around birth restores feeding in *Magel2* knockout (KO) mice^9^ and pediatric PWS patients^12^ suggesting predictive validity of this mouse model. Adult *Magel2* KO mice also present ASD-like features, such as social withdrawal ameliorated by perinatal OXT treatments^9,10^.

Modulation of social behavior with OXT has been at the center of many studies^13^ and it is now accepted that OXT contributes to filtering social salience signals^14^. OXT is used in numerous clinical trials^15^ with promising results for the treatment of ASD. While beneficial effects were observed in PWS patients treated with OXT, these effects are age dependent^16^ and not consistently found over different clinical trials^17^. Furthermore, chronic OXT treatments might be deleterious to specific aspects of social behavior^18,19^. This suggests that OXT therapy is not sufficient to treat social disabilities beyond the early postnatal critical period of neurodevelopment^20^ and stresses the need to better understand the OXT system and its modulators for treating social withdrawal in the adults.

Vasopressin (AVP), a peptide sharing many features with OXT^21^, is also important for the regulation of social salience and has been in the center of new clinical studies. ASD patients improved social communication upon treatment with AVP^22^ as well as with antagonists of AVP receptor subtype 1a (AVPR1a)^23^. The positive outcomes of both trials demands clarity about the mechanisms underlying AVPR responses in ASD and related diseases like PWS and SYS. We hypothesized that collective actions of OXT and AVP in the social salience brain network could explain the controversial efficacy of AVPR targeted therapies. Unfortunately, the roles of OXT and AVP have mainly been interrogated in isolation whereas combinatorial effects are anticipated^24^ given that both OXT and AVP are secreted in the brain upon social encounters^25^. Here, we devised strategies to understand the dual functions of AVP and OXT during social encounters in physiological conditions and in the context of *Magel2* deficiency.

## Results

### Social withdrawal linked to brain theta rhythmicity defects in *Magel2^+m/−p^* mice

Brain rhythmic activity, notably theta paced, across multiple regions of the social brain network is modulated by the novelty of social stimulus and associated with cognitive and emotional behaviors in humans and rodents^26,27^. Nonetheless, socially evoked theta rhythmicity has not been studied in animal models featuring autistic-like social withdrawal like the *Magel2^+m/−p^* mice. We used a telemetric system to record the brain electroencephalogram (EEG) from wire electrodes chronically implanted atop cortex as previously described^28^. Wired animals were subjected to multiple social trials with an unfamiliar juvenile (T1 to T4) followed by the encounter with a different juvenile (T5) to discriminate between degrees of novelty of the social stimulus (**Fig. 1a**). *Magel2^+m/−p^* mice performed poorly on the discrimination task with a mouse whereas object exploration was normal compared to WT littermate controls (**Fig. 1b**). At T1, EEG power spectral density analysis showed a socially induced modulation of activity in the theta band of healthy controls that is absent in *Magel2^+m/−p^* mice (**Fig. 1c**). The effect of *Magel2^+m/−p^* was specific of social stimulus as no difference with controls was observed during trials with objects (**Fig. 1c**). Changes of socially evoked theta rhythmicity correlated with the exploration time of conspecifics in WT controls but not in *Magel2^+m/−p^* mice (**Fig. 1d**). This contrasted with theta rhythmicity in trials with objects that did not correlate with the degree of novelty in WT controls but did in *Magel2^+m/−p^* mice (**Fig. 1d**). Therefore, this behavioral paradigm is sufficiently robust in *Magel2^+m/−p^* mice to detect social exploration disabilities consistent with brain theta rhythmicity defects particularly marked during the first social encounter.

**Figure 1.**
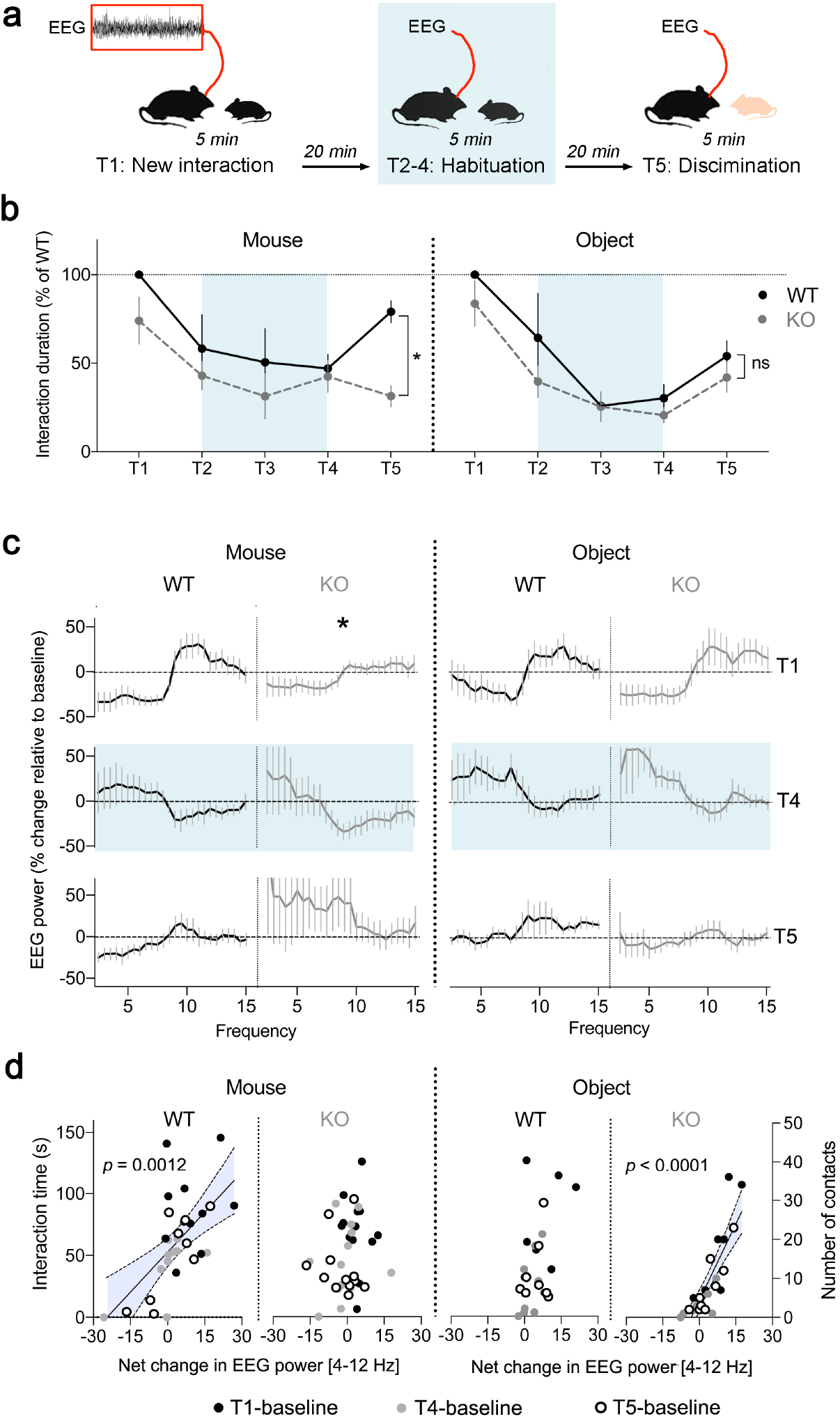
Deficits of EEG theta activity correlated with social defects in *Magel2^+m/−p^* mice. (**a**) Experimental timeline: EEG recorded during social habituation with a juvenile in 4 successive trials (T1-T4) of 5 min followed by a 5^th^ trial (T5) with a new conspecific. (**b**) Time exploring a mouse (left) or an object (right) throughout trials. Data (means±SEM) expressed as % of WT in n=15WT, 16KO mice for social and n=13WT, 15KO for non-social tests. Two-way ANOVA for social test: effect of trials *p*<0.0001, genotype *p*=0.028, and interaction *p*=0.035, post-hoc Sidak test comparing WT and KO at T5 \**p*<0.0001. For non-social test: effect of trials *p*<0.0001, genotype *p*=0.14, and interaction *p*=0.71. (**c**) Change in EEG power spectrum during trials. Data (means±SEM) expressed as % relative to baseline in n=11WT, 13KO mice for social and n=8WT, 8KO for non-social tests. Two-way ANOVA for social test: effect of genotype on the theta band at T1 *p*<0.0001, post-hoc Sidak test \**p*<0.05; at T4 *p*>0.9; at T5 *p*=0.009. For non-social test: no effect of genotype on the theta band. (**d**) Net changes of EEG power in the theta band during trials correlated with behavioral performance. For social test (interaction time in seconds): Spearman coefficient in WT *r*=0.56, *p*=0.0012 and KO *r*=0.21, *p*=0.18. For non-social test (number of contact with object): Spearman coefficient in WT *r*=0.39, *p*=0.053 and KO *r*=0.85, *p*<0.0001.

### Abnormal septal oxytocinergic and vasopressinergic systems in *Magel2^+m/−p^* mice

Previously, we showed that low number of OXTR binding sites specifically in the lateral septum (LS) of *Magel2^+m/−p^* mice covariate with social withdrawal^10^. This suggests a key role for the LS and its modulation by OXT in the physiopathology. C-Fos mapping showed more robust induction in the LS after interaction with a conspecific than with an object (**Fig. S1a**), an effect validated with p-S6 as indicator of rapid signaling (**Fig. S1b**). Compared to WT controls, basal expression of c-Fos was high and less reactive to social trials particularly in the LS dorsal (LSD) of *Magel2^+m/−p^* mice (**Fig. S1c,d**). Additionally, there were fewer and shorter AVP fibers in the LSD of *Magel2^+m/−p^* mice (**Fig. S2a,b**) as well as more OXT fibers in the septum (**Fig. S2c,d**). Such genotypic differences of topological innervations in septum suggest that both OXT and AVP actions may influence sociability between mice as previously hypothesized in healthy rats^29,30^.

To monitor the impact of AVP and the OXT analog agonist TGOT on theta rhythmicity, we combined bilateral intraseptal peptide injections with EEG recordings in freely moving cannulated mice (**Fig. 2a**). EEG power spectral analysis showed a trough of activity in the 4-8 Hz band and peak of activity in the 8-12 Hz band with AVP that contrasted with an opposite effect of TGOT (**Fig. 2b**). Such changes of theta rhythmicity mimicked patterns evoked in WT mice at social trials T1 and T4, respectively (*cf.* **Fig. 1c**). Implicitly, it suggests that a deficit of septal AVP at T1 might impair social exploration similar to the effect of *Magel2* deficiency. To test this possibility, we injected an AVPR antagonist, the Manning compound (MC) into the septum at T1 or T3 to assess its influence on social behavior and theta rhythm (see methods for details about *in vivo* pharmacology). Only at T1, the blockade of AVPR with MC impaired social exploration like the effect of *Magel2* deficiency (**Fig. 2c**). Similarly, and only at T3, the blockade of OXT receptors (OXTR) by intraseptal injection of the selective antagonist Atosiban impaired social exploration like the effect of *Magel2* deficiency (**Fig. 2c**). Consistently, the blockade of septal AVPR at T1 and OXTR at T3 modified theta rhythmicity throughout the remaining trials (**Fig. 2d**). Together, social exploration correlated with theta rhythmicity in these mice unless septal AVPR and OXTR were inhibited at T1 (**Fig. 2e**) and T3 (**Fig. 2f**), respectively. Therefore, timely actions of AVPR and OXTR in septum generate a sequence of brain activity patterns aligned with the degree of novelty of the social stimulus.

**Figure 2.**
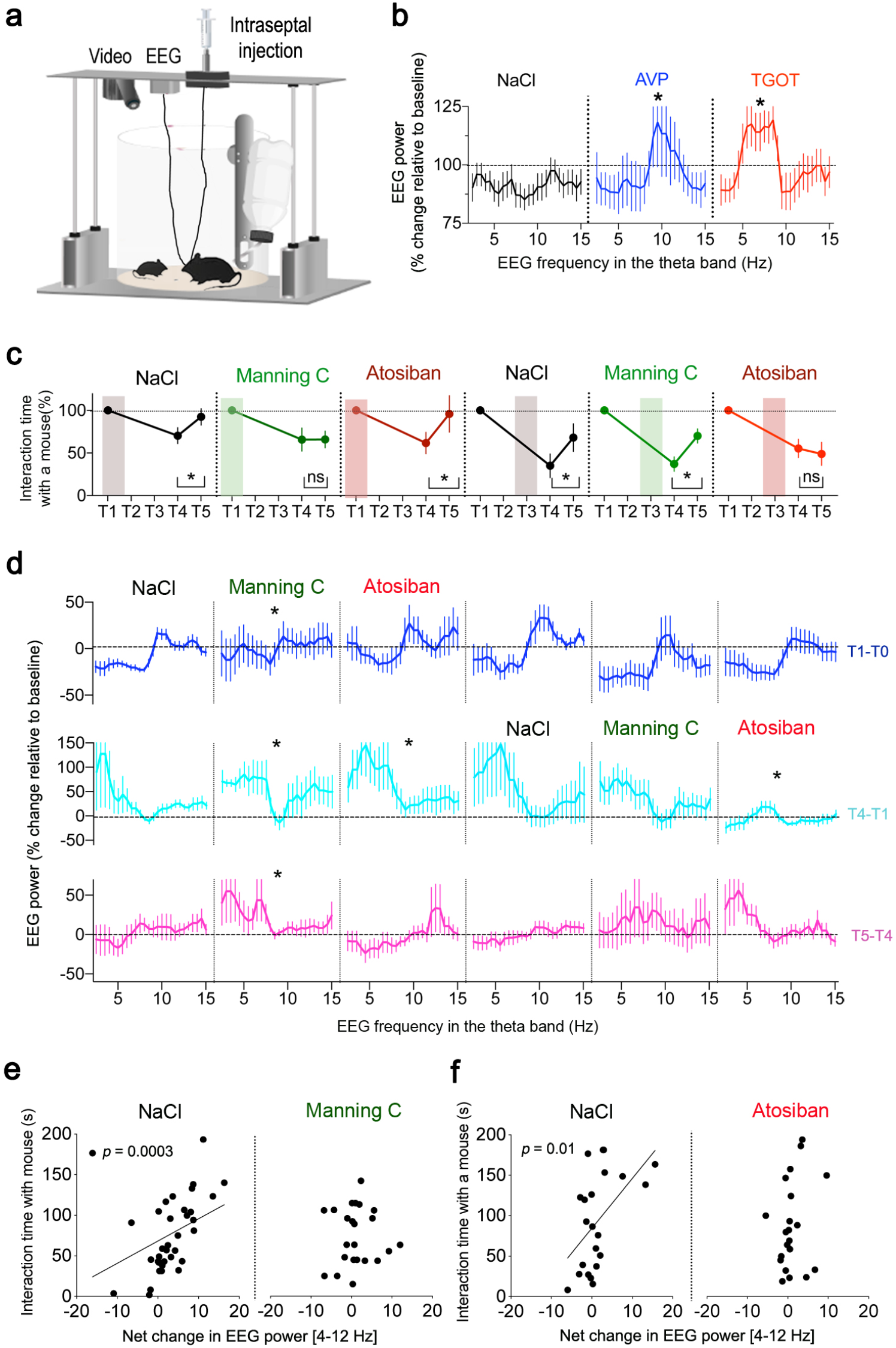
AVP and OXT receptors in septum modulate EEG theta rhythm during social behavior. (**a**) Experimental setup to inject drugs in cannulated septum while recording EEG in freely moving mice. (**b**) Percent change of EEG power in the low theta band [4-8 Hz] after intraseptal-injection of TGOT (3.10^−6^ M for 9 min) and in the high theta band [8-12 Hz] with AVP (3.10^−5^ M for 9 min). Means±SEM of n=13 NaCl, 10 AVP, 10 TGOT at 10 min post-injection: Kruskal Wallis test *p*=0.0011, effect of AVP \**p*=0.04 and TGOT \**p*=0.0005. (**c**) Time exploring a mouse throughout trials upon intra-septal injection of Manning Compound (10^−8^ M), atosiban (10^−8^ M) or vehicle at T1 or T3. Data (means±SEM) expressed as % relative to T1 in n=16 NaCL, 9 MC, 9 Atosiban mice injected at T1 and n=8 NaCL, 8 MC, 8 Atosiban mice injected at T3. Two-way ANOVA: effect of trials *p*<0.0001, post-hoc Dunnett test comparing T4 and T5 for NaCl \**p*=0.042, for Atosiban \**p*=0.023. Two-way ANOVA: effect of trials *p*<0.0001, post-hoc Dunnett test comparing T4 and T5 for NaCl \**p*=0.0017, for MC \**p*=0.0048. See the methods for the selectivity of MC and Atosiban on mOXTR and mAVPR. (**d**) Change of EEG power spectrum at T1 (top), T4 (middle) and T5 (bottom). Effect of intraseptal-injections at T1 of vehicle (NaCl), Manning C (MC 10^−8^ M) or atosiban (Ato 10^−8^ M). Data (means±SEM) expressed as % relative to baseline in n=11 NaCl, 8 MC, 7 Ato mice: Kruskal Wallis test at T1 *p*=0.021, effect of MC \**p*=0.0158; at T4 *p*=0.0015, effect of MC \**p*=0.0052 and Ato \**p*=0.0026; at T5 *p*<0.0001, effect of MC \**p*=0.007. Effect of intraseptal-injections at T3. Means±SEM of N=8 NaCl, 7 MC, 9 Ato mice: Kruskal Wallis test at T4 *p*<0.0001, effect of Ato \**p*<0.0001; at T5 *p*=0.0003, effect of Ato \**p*=0.0028. (**e**) Net changes of EEG power in the theta band between trials and baseline correlated with the interaction time with a mouse. Spearman coefficient if NaCl injected at T1: *r*=0.56, *p*=0.0003; if MC injected at T1: *r* = 0.03, *p*=0.86. (**f**) Net changes of EEG power in the theta band between trials and baseline correlated with the interaction time with a mouse. Spearman coefficient if NaCl injected at T3 *r*=0.53, *p*=0.01; if Ato injected at T3 *r*=0.3, *p*=0.17.

### Social salience depends on the AVP and OXT hypothalamoseptal circuits

To investigate timely activations of AVP and OXT neuronal networks, we first identified which cells responded throughout social trials to target their projections to LS with optogenetic constructs. We extended the initial c-Fos mapping to four brain regions containing AVP and OXT neurons: PVN, SON, BNST and LH^31–33^. AVP neurons were activated at T1 in the PVN of WT mice and in the BNST of *Magel2^+m/−p^* mice (**Fig. S3a,b**). OXT neurons were activated at T4 in the PVN of WT mice but less significantly in *Magel2^+m/−p^* mice (**Fig. S3c,d**). Thus, PVN neurons could be responsible for the release of AVP or OXT in the septum of WT mice unlike the septum of *Magel2^+m/−p^* mice that could rely on BNST neurons to secrete AVP.

To determine if the aforementioned sources of AVP and OXT modulate social behavior, we adopted an optogenetic silencing strategy. To this end, we used *Avp*-CRE and *Oxt*-CRE transgenic mice to independently target AAV virus coding for the CRE-dependent halorhodopsin (NpHR3.0-YFP) or eYFP into the PVN or BNST as indicated in **Fig. 3a**. CRE-mediated recombination was specific and efficient to express NpHR3.0-YFP either in OXT neurons of PVN or in AVP neurons of PVN or BNST (**Fig. 3b**). A prerequisite to operate as PVN-LS or BNST-LS circuits responding to social trials was that these neurons projected YFP-positive axon boutons into the septum (**Fig. S4a**). As expected, yellow light stimulation of recombinant NpHR3.0 in PVN of coronal brain slices reduced the firing rate of target neurons (**Fig. 3c**). In LS coronal slices, patch clamp recordings of predefined AVP-responding cells or TGOT-responding cells determined the impact of optogenetic manipulations specifically at axon boutons (**Fig. S4b**). Yellow light stimulation of NpHR3.0 on these cells had no effect in absence of social trials (**Fig. S4b-d**). On the contrary, blue light stimulation of ChR2-YFP (expressed with similar viral strategy) evoked responses typical of AVP in predefined AVP-responding cells in *Avp*-CRE animals (**Fig. S4e**) as well as TGOT in predefined TGOT-responding cells in *Oxt*-CRE animals (**Fig. S4f**). Importantly, blue-light evoked responses were blocked by the AVPR antagonist at AVP-responding cells, and OXTR antagonist at TGOT-responding cells. These results validated the optogenetic control of OXT or AVP releases from axon boutons in LS.

**Figure 3.**
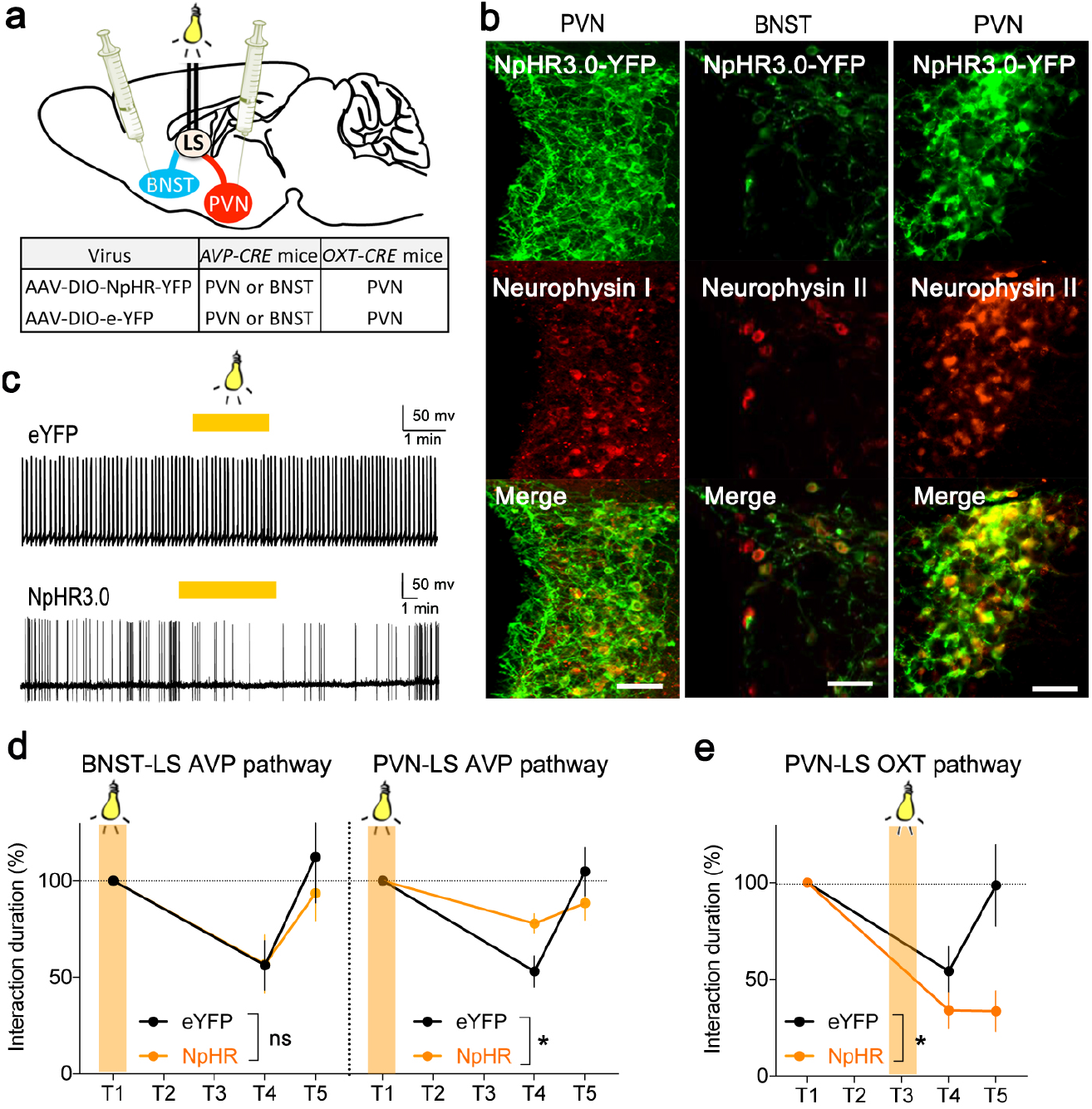
Modulation of social behavior by optogenetic inhibition of the OXT and AVP septohypothalamic circuit pathways. (**a**) Viral-mediated optogenetic silencing of septal inputs from OXT neurons (*Oxt*-CRE mice) in hypothalamic paraventricular nucleus (PVN) or from AVP neurons (*Avp*-CRE mice) in PVN or Bed nucleus of stria terminalis (BNST). (**b**) Co-expression of CRE-mediated NpHR3.0-YFP with Neurophysin I (OXT) in PVN neurons or with Neurophysin II (AVP) in PVN as well as BNST neurons. Scale=25 μm. (**c**) Firing of action potentials recorded in whole cell configuration in OXT neurons expressing NpHR3.0-YFP or eYFP. Stimulation of cell body with yellow light decreased firing rate. (d) Time exploring a mouse throughout trials. Data (means±SEM) expressed as % relative to T1 in each group of n=8 eYFP, 9 NpHR3.0 in BNST and 11 eYFP, 10 NpHR3.0 in PVN of *Avp-CRE* mice. Two-way ANOVA: Interaction of trials and NpHR3.0 stimulation with light (561nm, continuous stimulation, 5min, ~2mW at T1) of BNST fibers: *p*=0.6; of PVN fibers: *p*=0.01 post-hoc Sidak test comparing eYFP with NpHR3.0 at T4 \**p*=0.02. (**e**) Time exploring a mouse throughout trials. Data (means ± SEM) expressed as % of T1 in each group of n=11 eYFP, 7 NpHR3.0 in PVN of *Oxt-CRE* mice. Two-way ANOVA: Interaction of NpHR3.0 stimulation of PVN fibers with light (561nm, continuous stimulation, 5min, ~2mW at T3) and trials: *p*=0.05, post-hoc Sidak test comparing eYFP with NpHR3.0 at T5 \**p*=0.015.

Optic fibers were chronically implanted atop LS bilaterally of WT mice to achieve light-dependent silencing of projecting axons from AVP neurons at T1 and from OXT neurons at T3. We found that yellow light stimulation of NpHR3.0 in the PVN-LS AVP pathway (**Fig. 3d** right) and the PVN-LS OXT pathway (**Fig. 3e**) impaired social exploration distinctly. Hypothalamoseptal pathways activated timely to generate a functional sequence of AVP and OXT septal releases according to the degree of novelty of the social stimulus. In contrast, silencing of an extra-hypothalamoseptal pathway, the BNST-LS pathway failed to modify exploration through social trials (**Fig. 3d** left), highlighting remarkable specificity about AVP input source to the LS of WT mice under these behavioral conditions.

### Disarray of AVP and OXT septal releases disrupted exploration of social salient stimuli

As sequential septal release of AVP and OXT is critical to express social exploration, we aimed to disrupt this orderly sequence during social trials with optogenetic stimulation of the hypothalamoseptal pathways. We used CRE-dependent ChR2-YFP or eYFP constructs delivered into the PVN or BNST of *Avp*-CRE and *Oxt*-CRE mice and induced light stimulation of LS projecting axons from the BNST-LS or PVN-LS circuits to alter the orderly sequence of AVP and OXT releases (**Fig. S5a-c**). Deficits of social exploration manifested if AVP was released at T3 instead of T1 from BNST-LS pathway and if OXT was secreted at T1 instead of T3 from PVN-LS pathway (**Fig. S5d-e**). Therefore, it is not the releases of AVP and OXT per se, but its orderly sequence aligned to the degree of social novelty that determined social exploration.

### Paucity of cells responding to AVP and OXT orderly sequence in LS of *Magel2^+m/−p^* mice

Cellular targets in the LS of AVP and OXT orderly sequence remained to be explored. We used patch clamp recordings in coronal brain slices to characterize neurons in LS based on changes of firing rate upon bath application of AVP or TGOT. Half of the cells were selectively excited by AVP (type I) whereas the others were either stimulated selectively by TGOT (type II), or inhibited by both peptides (type III) (**Fig. 4a**). Retrobead anatomical tracing (**Fig. 4b**) revealed that the type II and III cells mostly, projected to the medial septum (MS). Connection between this pathway and the hippocampus^34^ is known to organize sequential content of sensory experiences via theta-paced sequence of cell assemblies^35–37^. Modulation of sequential content by LS neurons may rely on inputs containing AVP from the hypothalamus, and glutamate from hippocampus^38^, whereas outputs to MS are enriched with OXTR, suggesting that both peptides could act at different levels of this circuit. Electrophysiological response of the type III cells depended on the orderly sequence of AVP and TGOT contrary to the other cell types recorded. That is, AVP must be presented first to gain responsiveness to TGOT while the effect of AVP was unconditional to the order of presentation (**Fig. 4c**). These neurons also differed in terms of spontaneous activity patterns, morphology, and other electrophysiological properties (**Fig S6**).

**Figure 4.**
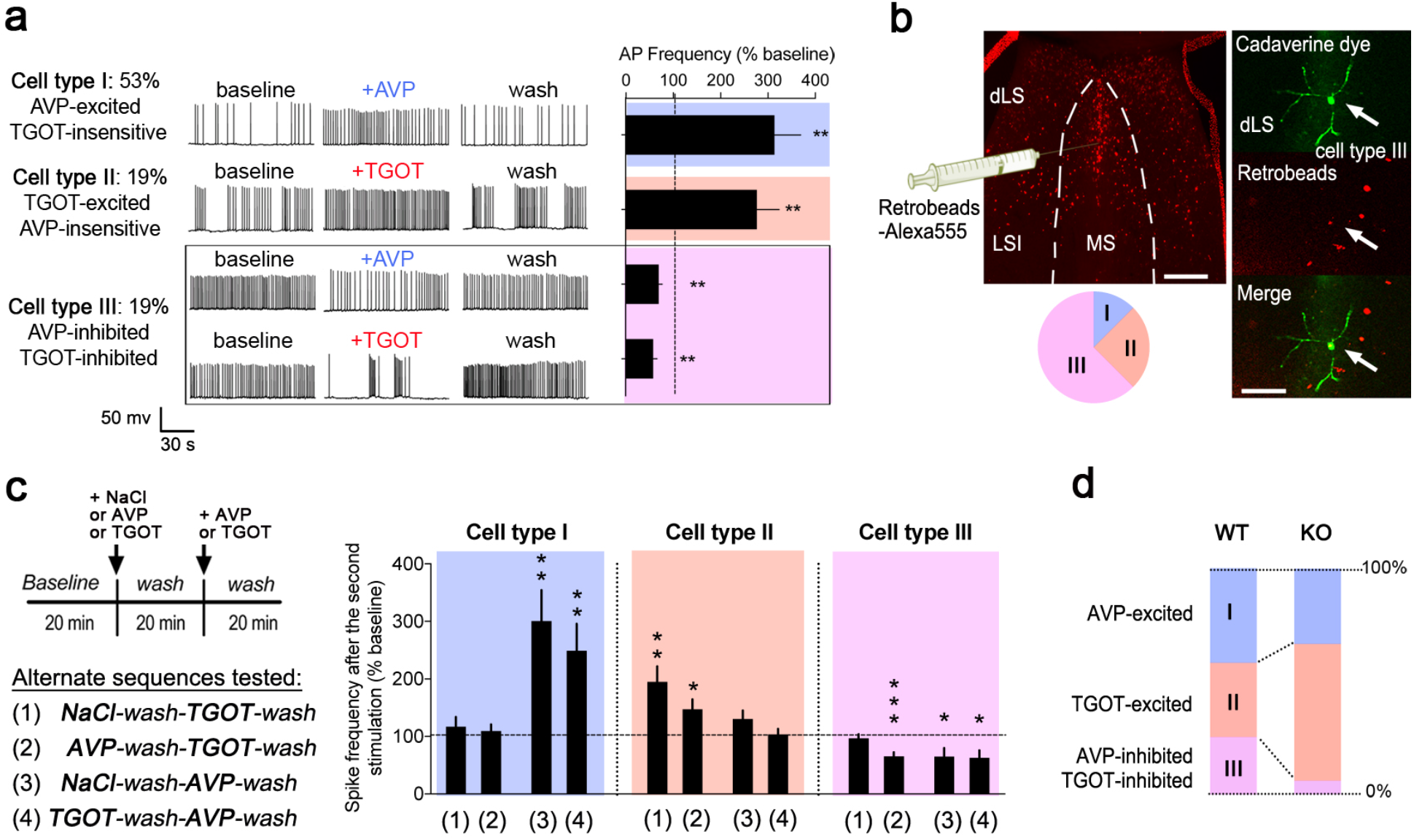
Paucity of cells responding to the orderly sequence of AVP and OXT in LS of *Magel2^+m/−p^* mice. (a) Firing rate of action potentials recorded in whole cell configuration in septal slices at baseline and upon 2-min bath application of AVP (10^−6^ M) and TGOT (10^−7^ M). All 3 types of response reverted to baseline after washout. Data (means±SEM) expressed as % relative to baseline prior treatment in n=29 TGOT-excited, 34 AVP-excited, 20 AVP-inhibited and 16 TGOT-inhibited cells. T-Test for the effect of TGOT-excitation \*\**p*=0.0007; AVP-excitation \*\**p*=0.0005; TGOT-inhibition \*\**p*<0.0001; AVP-inhibition \*\**p*=0.001. (**b**) Cells in LSD marked with cadaverine-Alexa-Fluor-594 via the patch pipette corresponded mostly to the type III if they uptake fluorescent retrobeads injected in MS (n=16 neurons, 2 type I, 4 types II, 10 type III in 11 mice). Scales=200 and 50 μm. (**c**) Effect of orderly sequence of AVP and TGOT on spike frequency. Sequence of 2 stimuli (2 min each) to categorize cell types *a posteriori*. Wilcoxon test for the effect of TGOT alone on type II cells \*\**p*=0.0027; AVP 1^st^–TGOT 2^nd^ on type II cells \**p*=0.0155; AVP 1^st^–TGOT 2^nd^ on type III cells \*\*\**p*<0.0001; AVP alone on type I cells \*\**p*=0.0018; TGOT 1^st^-AVP 2^nd^ on type I cells \*\**p*=0.0055; AVP alone on type III cells \**p*=0.039; TGOT 1^st^-AVP 2^nd^ on type I cells \**p*=0.0138. (**d**) Proportion of cells categorized as a function of their responses to AVP and TGOT on the frequency of action potentials. n=28WT, 11KO type I cells, 22WT, 20KO type II cells and 17WT, 2KO type III cells in septal slices of 11 *Magel2^+m/−p^* and 29 WT controls mice.

Importantly, type III neurons were scarcer while type II neurons were denser in *Magel2^+m/−p^* mice than in WT controls (**Fig. 4d**). Such a change of proportion between cell types could depend on OXTR and AVPR dual expression. To distinguish AVPR and OXTR binding sites with cellular resolution, we synthetized d [Lys (Alexa-Fluor-647)^8^] VP, a fluorescent peptide selective for mouse OXTR *in vitro* (**Fig. 5a**) and *in vivo* if co-injected with the competitive AVPR ligand MC (**Fig. 5b** bottom left). For a rather selective labeling of mouse AVPR (**s**) *in vivo*, higher dose of the fluorescent peptide was used with the competitive OXTR ligand TGOT (**Fig. 5b** bottom right). When injected in LS after social trials, d [Lys (Alexa-Fluor-647)^8^] VP marked cells equipped with OXTR or AVPR, some of which also contained the activity-dependent indicator p-S6 and retrobeads (**Fig. 5c**). Specifically, LS cells projecting to MS were more abundant and more responsive to social trials in WT controls than in *Magel2^+m/−p^* mice (**Fig. 5d,e**). These cells are mostly GABAergic somatostatin neurons labelled with retrobeads (likely the type III, **Fig. S7**). All in all, the septum of *Magel2^+m/−p^* mice is ill equipped to organize sequential content of social signals evoked by AVP and OXT releases (**Fig. 5f**).

**Figure 5.**
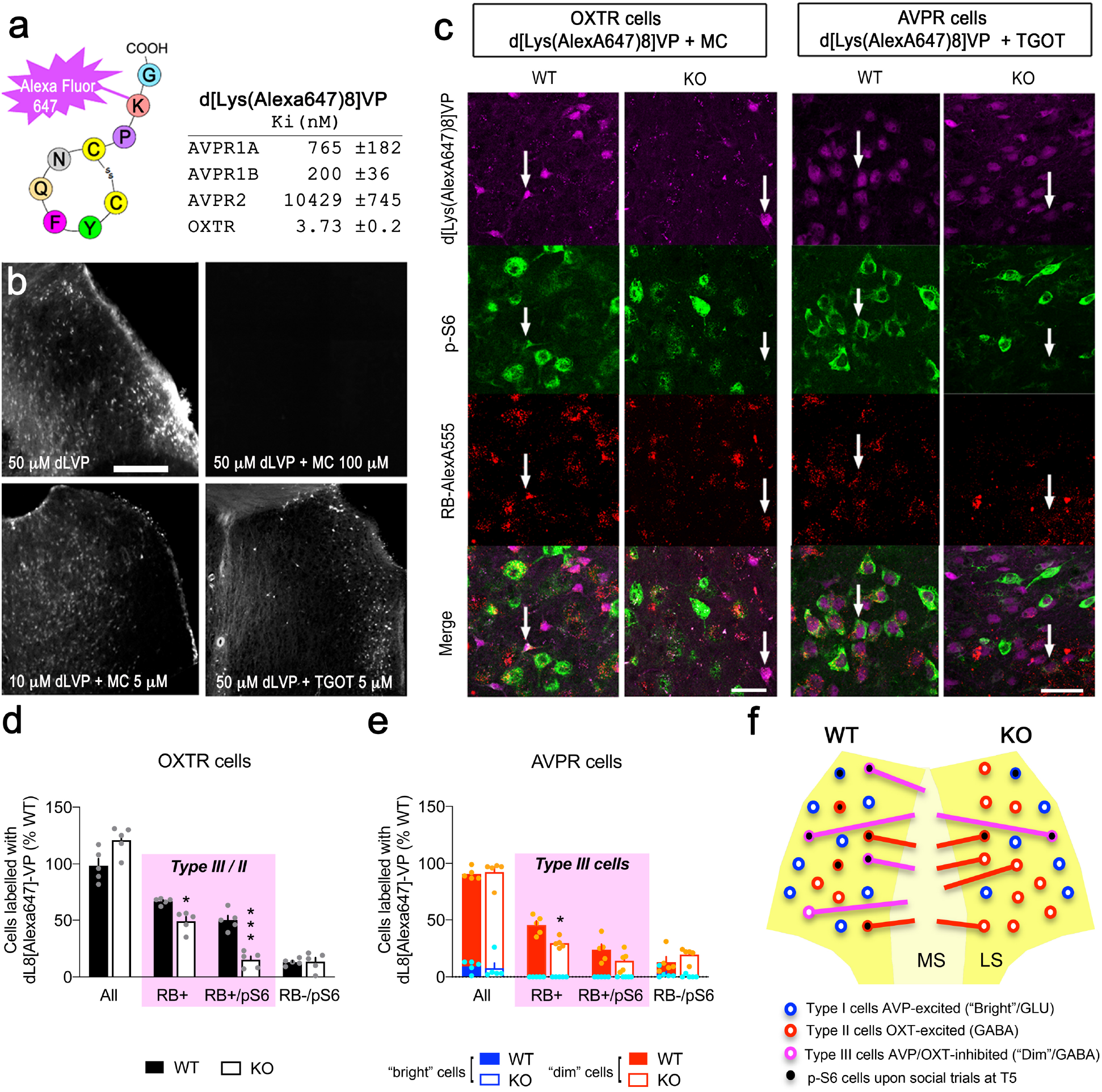
A new fluorescent ligand to identify OXT and AVP responding cells *in vivo*. (**a**) Affinity of d[Lys(Alexa-Fluor-647)^8^]VP *in vitro* for the indicated mouse receptors: AVPR1b and OXTR transfected in HEK293 cells and AVPR1a and AVPR2 endogenous from liver and kidney, respectively. Means±SEM of n=3 independent competition assays against [^3^H]AVP. (**b**) Representative *in vivo* labeling of cells with intraseptal injection of d[Lys(Alexa-Fluor-647)^8^]VP in control mice competed entirely when co-injected with the non selective OXTR/AVPR antagonist (100μM MC), and partially with 5μM MC (to block AVPR sites only) or with 5μM TGOT (to block OXTR sites only). Scale=200μm. See the methods for the selectivity of MC and TGOT on mOXTR and mAVPR. (**c**) Representative *in vivo* labeling of OXTR (left) and AVPR (right) binding sites in LS of WT and *Magel2^+m/−p^* mice prior injected with retrobeads (RB-Alexa-Fluor-555) in MS to mark cell type III. Cannulated animals were administered with drugs in LS 5min after the last trial of social habitation/discrimination and 10min later, sacrificed. Scale bars=25μm. (**d**) Percent of cells expressing OXTR binding sites and p-S6 in LS. Total OXTR cells 353WT of which 237RB+, 451KO of which 224RB+. Means±SEM of n=5 WT and 5 *Magel2^+m/−p^* mice. Two-way ANOVA: Effect of genotype on OXTR cells: *p*<0.0001, post-hoc Sidak test for the percent of OXTR cells projecting to MS \**p*=0.0415 and OXTR cells projecting to MS with pS6 \*\*\**p*=0.0002. (**e**) Percent of cells expressing AVPR binding sites and p-S6 in LS. Total AVPR cells 474WT of which 235RB+, 379KO of which 112RB+. Means ± SEM of n=5 WT and 5 *Magel2^+m/−p^* mice. Two-way ANOVA: Effect of genotype on AVPR cells: *p*=0.0256, post-hoc Sidak test for the percent of AVPR cells projecting to MS \**p*=0.011. (**f**) Summary of cell types equipped with AVPR/ OXTR in LS. Type III cells inhibited by both AVP and TGOT are less solicited (p-S6 signaling) in *Magel2^+m/−p^* mice compared to WT controls during social trials.

### AVPR priming in LS of *Magel2^+m/−p^* mice restored exploration of social salient stimuli

To normalize theta-paced sequence of cell assemblies in the LS of *Magel2^+m/−p^* mice, we promoted AVPR septal response during the first social encounter (**Fig. S8d**). For this, *Magel2^+m/−p^* mice were cannulated in LS to receive bilateral AVP injections at T1, which increased theta rhythmicity throughout trials (**Fig. S8a**) and restored social exploration (**Fig. 6a**). These activities were not correlated if NaCl or AVP+Atosiban were injected instead of AVP alone (**Fig. 6b** and methods for details about *in vivo* pharmacology). Consistently, NaCl or AVP+Atosiban injections failed to restore theta rhythmicity and social exploration (**Fig. 6a**, **S8a**). Thus, inhibition of septal OXTR with Atosiban despite AVPR priming highlighted the necessity of AVPR and OXTR collective responses to restore social behavior of *Magel2^+m/−p^* mice.

**Figure 6.**
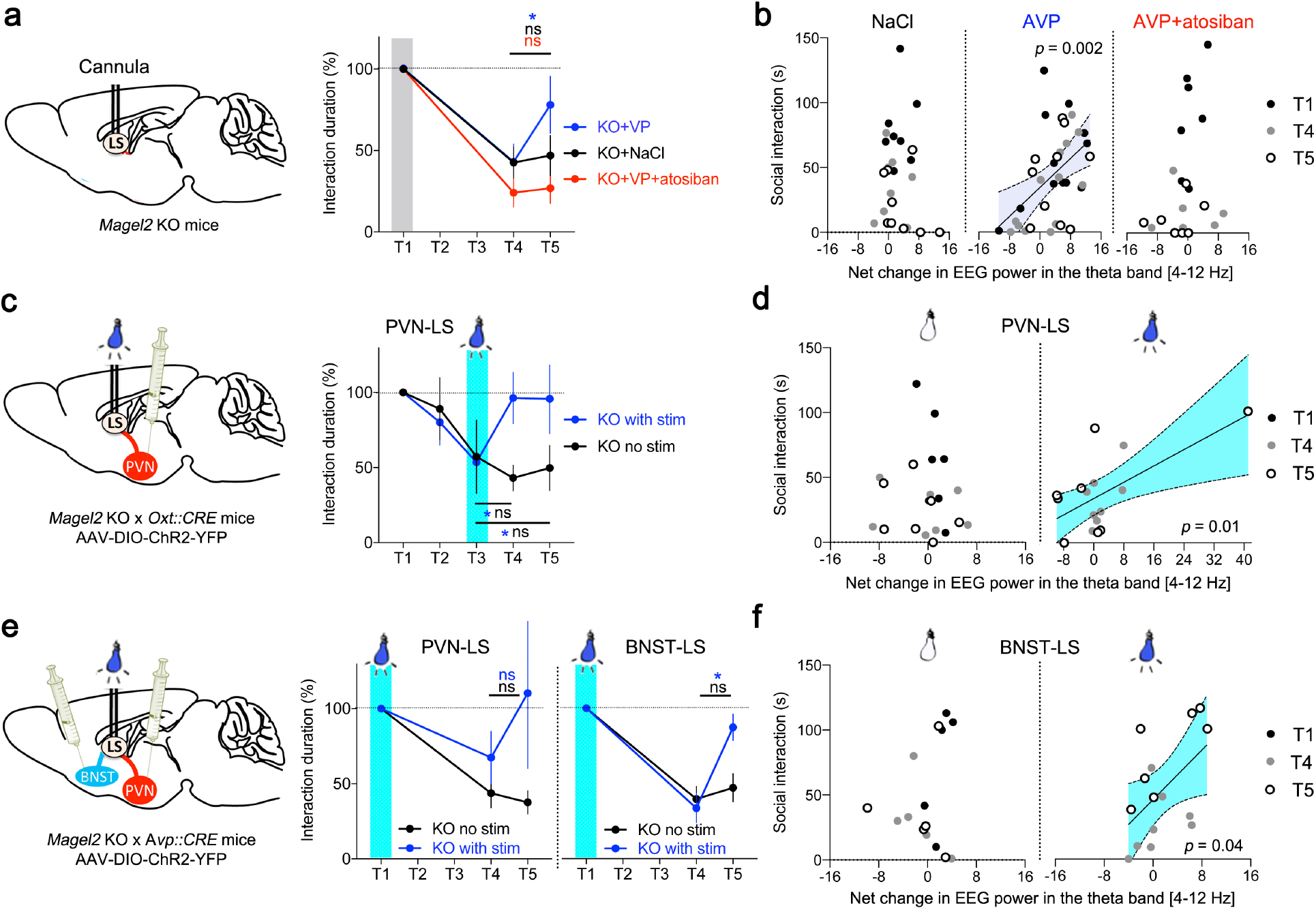
Stimulation of the BNST-LS AVP pathway and PVN-LS OXT pathway restored social behavior in *Magel2^+m/−p^* mice. (**a**) Time *Magel2^+m/−p^* mice explored WT conspecifics after intraseptal injection of AVP (3.10^−6^ M) with or without atosiban (5.10^−8^ M) at T1. Data (means±SEM) expressed as % relative to T1 in each group of n=13NaCl, 12AVP and 6AVP+atosiban mice. Two-way ANOVA: Effect of social trials: *p*<0.0001 post-hoc Dunnett test for the effect of NaCl: T4 vs T1 \**p*<0.0001; effect of AVP: T4 vs T1 \**p*<0.0001 and T4 vs T5 \**p*=0.01; effect of AVP+atosiban: T4 vs T1 \**p*<0.0001. (**b**) Net changes of EEG power in the theta band between trials correlated with time of social interaction. Pearson coefficient if NaCl injected at T1: *r*=−0.08, *p*=0.6; if AVP injected at T1: *r*=0.49, \**p*=0.024; if AVP+atosiban injected at T1: *r*=0.3, *p=*0.18. (**c**) Time *Magel2^+m/−p^* mice explored WT conspecifics after blue light stimulation (473nm, 30Hz, 10ms pulses for 2min, 2mW) of OXT neuron projections between PVN and LS (PVN-LS pathway). Data (means±SEM) expressed as % relative to T1 in each group of n=12 no stim, 12 stim mice. Two-way ANOVA: Effect of ChR2 stim on social trials: *p*=0.0037 post-hoc Dunnett test for the effect of no stim: T3 vs T1 \**p*=0.035; effect of ChR2 stim: T3 vs T1 \**p*=0.016 T3 vs T4 \**p*=0.029 and T3 vs T5 \**p*=0.0318. (**d**) Net changes of EEG power in the theta band between trials correlated with time of social interaction after optogenetic stimulation of PVN-LS OXT terminals. Pearson coefficient if no stim at T3: *r*=−0.2, *p*=0.47; if ChR2 stim at T3: *r*=0.62, \**p*=0.01. (**e**) Time *Magel2^+m/−p^* mice explored WT conspecifics after blue light stimulation (473nm, 20Hz, 5ms pulses for 2min, 2mW) of AVP neuron projections between BNST and LS (BNST-LS pathway) or PVN and LS (PVN-LS pathway). Data (means±SEM) expressed as % relative to T1 in each group of n=9 no stim, 6 stim BNST-LS mice and 9 no stim, 9 stim PVN-LS mice. (Left) Two-way ANOVA: Effect of ChR2 in PVN-LS on social trials: *p*=0.2 post-hoc Dunnett test for the effect of no stim: T4 vs T1 \**p*=0.0008 and T4 vs T5 *p*=0.75; effect of stim: T4 vs T1 *p*=0.16 and T4 vs T5 *p*=0.6. (Right) Two-way ANOVA: Effect of ChR2 in BSNT-LS on social trials: *p*=0.005 post-hoc Dunnett test for the effect of no stim: T4 vs T1 \**p*<0.0001 and T4 vs T5 *p*=0.6; effect of stim: T4 vs T1 \**p*<0.0001 and T4 vs T5 \**p*=0.01. (**f**) Net changes of EEG power in the theta band between trials correlated with time of social interaction. Pearson coefficient if no stim at T1: *r*=−0.17, *p*=0.07; if ChR2 stim at T1: *r*=0.53, \**p*=0.04.

In a second experiment, we promoted the OXT system of *Magel2^+m/−p^* mice given its clinical potential for alleviating social disabilities in humans^39,40^. To this end, we used optogenetic stimulation of CRE-dependent ChR2-YFP recombined in OXT neurons of PVN in *Oxt*-CRE x *Magel2^+m/−p^* mice to promote OXT septal release during social habituation. This manipulation increased exploration duration with known and unknown mice without discrimination (**Fig. 6c**), and exploration duration correlated with changes of theta rhythmicity (**Fig. 6d**). Theta rhythm of *Magel2^+m/−p^* mice optogenetically-stimulated for OXT septal release (**Fig. S8b**) looks alike that of WT mice optogenetically-deprived of AVPR septal response at T1 (**Fig. 2d**). Thus, OXT therapies may not be optimally effective in diseases characterized by AVPR sequential priming defects.

In a third experiment, we used optogenetic stimulation of ChR2-YFP recombined in AVP neurons of PVN or BNST of *Avp*-CRE x *Magel2^+m/−p^* mice to simulate AVP release normally evoked by the first social encounter. We found that blue light-stimulation of the BNST-LS AVP pathway promoted consistent social exploration (**Fig. 6e** right) unlike stimulation of the PVN-LS AVP pathway (**Fig. 6e** left). Optogenetic stimulation of BNST-LS AVP terminals of *Magel2^+m/−p^* mice modulated theta rhythmicity (**Fig. S8c**) that correlated with social exploration (**Fig. 6f**). This result indicates that social behavior is restored in *Magel2^+m/−p^* mice by promoting AVPR septal response during the first encounter through the BNST-LS AVP pathway.

Collectively, one promising avenue for disease modification is to restore the orderly sequence of AVP and OXT responses for organizing theta-paced sequence of social salient information flow through the septum.

## Discussion

One major feature of physiopathology associated with *Magel2* deficiency reported in this study is the abnormal functional topology of both the AVP and OXT neuronal networks innervating the septum. This is decisive because AVP and OXT must act collectively on septal neurons to demonstrate social salience. Normally, AVP acts first on septal neurons upon release evoked by the novelty of the social stimulus while OXT acts second on septal neurons upon release elicited by the repetition of the social stimulus as previously suggested in rats^41,42^. It is the degree of social novelty that commanded sequential activations of AVP and OXT hypothalamoseptal neuronal networks as seen with the c-Fos mapping studies.

Pathologically, not only septal AVP fibers were scarcer and PVN AVP neurons poorly activated upon social novelty but also a cryptic extra-hypothalamoseptal AVP network originating from the BNST was activated by social encounters. This suggested that a functional map-to-action relevant for expressing sequential content as described in the hippocampus^36^ could differ between *Magel2^+m/−p^* mice and WT controls for perceiving the degree of social novelty. Perhaps, new social encounters are seen as threatening more than rewarding by *Magel2^+m/−p^* mice, resulting in the mobilization of alternate circuit pathways that influence behavioral response. For instance, threats, unpredictability and social anxiety activate the BNST in which the lesion of AVP cells specifically, reduced social anxiety and aggressiveness^43,44^. Many brain regions among which the PVN and BNST induced *c-Fos* and either *Oxtr* or *Avpr(s)* if conspecific odors came from healthy individuals or sick individuals, respectively. Further blockade of AVPR inhibited social avoidance to sick odors^45^, thus providing evidence that socially evoked activation of AVP BNST cells is an appropriate response to threats that *Magel2^+m/−p^* mice may privilege even with healthy conspecific. In fact, activation of LSD neurons, a region rich of AVP fibers, inhibits aggressive behavior in mice via its projections to the ventromedial hypothalamus^46^. In *Magel2^+m/−p^* mice, c-Fos activation was elevated in all septal areas under isolation, failing to respond upon social encounters unlike healthy controls, indicating a possible conflict between social perception and theta paced neuronal activation in the social brain network^27^. Despite their social disabilities, *Magel2^+m/−p^* mice perform well with object exploration, exhibiting correlated activities with theta rhythmicity. This contrasts with the WT mice that did not show such correlated activities in the object trials. In agreement, the sensitivity for discriminating faces and objects was reported respectively, impaired and enhanced in adolescents with ASD^47^ which adds to the face validity of the *Magel2^+m/−p^* mice as model of ASD.

Abnormalities in the OXT PVN-LS pathway were less prominent than in the AVP system of *Magel2^+m/−p^* mice. This is surprising considering the alteration of oxytocinergic neurons function described in *Magel2^+m/−p^* mice^48^ but could be due to compensatory mechanisms on post-synaptic target cells in the LSD such as the replacement of type III neurons by the type II or such as the increase of OXT fibers in the LS. Besides the apparent disarray between AVP and OXT septal releases in *Magel2^+m/−p^* mice, abnormal septal response to social salience hormones corresponded to the underrepresentation of type III neurons belonging to the LS-MS pathway previously described as sequence generator linked with the hippocampus^37,49^. These neurons, which are unconditionally inhibited by AVPR, must gain competence to be inhibited by OXTR thereby operating as detectors of coincidence organizing the orderly sequence of AVP and OXT septal response within a functional map-to-action. Despite the septum of *Magel2^+m/−p^* mice is ill equipped, promoting AVPR septal response during social novelty restored social behavior more efficiently than by stimulation of OXTR septal response during social habituation. Therefore, restoring a complete orderly sequence of AVPR and OXTR septal releases is essential for organizing sequential content of social salience signals. This mechanism could also have implications for treating human pathologies given that activity of the hypothalamoseptal areas was associated with affiliative emotion^50^, and that OXT given intranasally increases the functional connectivity between the septum and other key areas of the social salience and reward circuits^51^.

Therapeutic priming of AVPR septal response for a few minutes at the time of social novelty restored social discrimination more than 1h later by a mechanism requiring OXTR septal response in *Magel2^+m/−p^* mice. Molecular and electrophysiological studies provided some clues to understand this effect. First, AVP-deficient Bratteleboro rats centrally administered with AVP corrected for several hours the frequency deficit of theta rhythm^52^. Second, AVPR and OXTR responses overlap in the septum where theta paced sequence of cells assemblies projecting to the hippocampus are modulated by OXT and AVP^53^. Third, sequential content of AVP and OXT in septum is suspected to modify theta paced network activity during social encounter. Consistent with an AVPR priming effect in the LS, AVP stimulation of LS neurons was previously shown to condition subsequent excitatory response to glutamatergic inputs from the hippocampus in rats^54,55^. Fourth, cells detecting the coincidence of AVP and OXT sequential content are underrepresented in septum of *Magel2^+m/−p^* mice. They are GABAergic somatostatin-positive cells (type III) projecting to MS likely equipped with both AVPR and OXTR. Unfortunately, this remains an open question, as d [Lys (Alexa-Fluor-647)^8^] VP did not allow for co-labeling of OXTR and AVPR. Lack of AVPR priming during social novelty impaired theta paced information flow through the septum to express social salience by OXTR modulation during habituation. So, therapeutic AVPR priming of *Magel2^+m/−p^* mice should restore septal network activity in response to social encounter. Consistently, intranasal administration of AVP (but not OXT) increased reciprocated collaboration between humans and its associated reactivity in the LS^56^ even several days beyond treatment^57^.

In terms of clinical perspectives, it is encouraging that priming of septal AVPR can be achieved even by a cryptic source of AVP (*e.g.* the BNST-LS pathway), further illustrating that circuit defects can be alleviated by loading the septum with AVP at the right time. Moreover, theta paced septal activity and its modulation by social salience hormones is an opportunity to use EEG recordings for predicting social behavior outcomes as demonstrated in this study and others^27^. In humans, EEG abnormalities and epilepsy have been reported in patients with PWS^58^ and ASD^59^. Few studies reported deficits of social task-related EEG power spectrum changes in ASD patients^60,61^. Here, we provide not only an EEG signature of social disabilities in *Magel2^+m/−p^* mice but also a blueprint of traces specific for AVPR and OXTR modulations in mouse septum corroborated in rats^53^. Future research will focus on improving an EEG predictive marker of the sensitivity to AVP and OXT in related pathologies and therapies. This is particularly relevant in pathologies such as PWS, SYS and ASD because of the heterogeneity of clinical features and the responses to treatments (*e.g.* bumetamide^62^ and oxytocin^63^).

## CONTRIBUTIONS

A.M.B, M.G.D and F.J designed and verified analytical methods. A.M.B, Y.D, D.D and D.H carried behavior studies. A.M.B, M.G.D carried electrophysiological studies. A.M.B and Y.D carried stereotaxic injections. Cs.T, A.O and M.M synthesized d[Lys(Alexa-Fluor-647)^8^]VP, characterized *in vitro* by G.G and *in vivo* by F.J and Y.D. E.P performed histology. D.D and E.P verified implantations in postmortem brains. A.M.B and D.D analyzed EEG. F.M and P.C provided critical feedback. A.M.B and F.J wrote the manuscript. All authors reviewed and approved the final manuscript. The authors declare no competing financial interests.

## ACKNOWLEDGEMENTS

This work is supported by ANR (M.G.D, F.M), Fondation Lejeune (M.G.D), Fondation pour la recherche médicale (F.J, A.M.B), Centre hospitalier de Montpellier (D.D., P.C.), Montpellier University (A.M.B), and generous support from R. Makineni, R. Tyner, F. Paulsen (M.M.). We thank from IGF in Montpellier, N. Marchi for sharing EEG devices, B. Boussadia for advices on EEG, M. Arango-Lievano for technical strategies, critical reading of the manuscript and M. Tauber (CHU Toulouse) for discussions on PWS.

## ABBREVIATIONS

AAV: Adeno-associated virus;
ASD: Autism spectrum disorder;
Ato: Atosiban;
AVP: Arginine-Vasopressin;
AVPR: AVP receptor;
BNST: Bed stria terminalis nucleus;
ChR2: Channel rhodopsin-2;
EEG: electroencephalogram;
GABA: gamma-aminobutyric acid;
GLU: glutamate;
KO: Knockout;
LH: Lateral hypothalamus;
LS: Lateral septum;
LSD: LS dorsal;
LSI: LS intermediate;
LSV: LS ventral;
Magel2: MAGE family member L2;
MC: Manning Compound;
MS: Medial septum;
NpHR3: Halorhodopsin-3;
OXT: Oxytocin;
OXTR: OXT receptor;
PVN: Paraventricular nucleus;
PWS: Prader-Willi Syndrome;
p-S6: phospho-protein ribosomal S6;
SON: Supraoptic nucleus;
SYS: Schaaf-Yang syndrome;
WT: wildtype;
YFP: Yellow protein fluorescent.

## Methods

All experiments were carried out in accordance with the Directive by the Council of the European Communities (86/609/EEC) following guidelines from the French Ministry of research and ethics committee for the care and use of laboratory animals (approved protocol APAFIS 5133, 8940, 11468).

### Reagents

Vasopressin, TGOT, Atosiban, Manning Compound, TTX, CGP35348, GABAzine and salts for electrophysiology are from Sigma Aldrich (S.A.R.L, Saint-Quentin, France). Antibodies: mouse monoclonal Neurogranin (Ab5620), NeuN, GAD67, Calretinin (Merck Millipore, St Quentin en Yvelines, France), Calbindin (Swant, Marly, SWI), Guinea Pig polyclonal Neurotensin (Synaptic systems), rat polyclonal Somatostatin, chicken polyclonal GFP (Abcam, Cambridge, UK), rabbit polyclonal c-Fos, p-S6 (Cell Signaling Technology, Ozyme France) and custom-made rabbit polyclonal antibodies against neurophysin I and II (Kindly provided by H. Gainer at NIH, Bethesda, USA).

### Animals

Oxttm1.1(CRE)Dolsn/J (*Oxt*-CRE) and Avptm1.1(cre)Hze/J (*Avp*-CRE) male mice were purchased (Jackson labs, Bar Harbor, MA, USA) and maintained on a C57B6/J background (Janvier Labs, Le Genest-Saint-Isle, France). *Magel2*-deficient mice were provided by F. Muscatelli (INMED, Marseille, France) and backcrossed with *Oxt*-CRE and *Avp*-CRE lines on a C57B6/J background. *Magel2*^+m/−p^ (paternal KO allele) mice were obtained by crossing WT females with ^+p/-m^ (maternal KO allele) males. *Magel2* being a maternally imprinted gene, *Magel2*^+m/−p^ mice did not express *Magel2* and were considered functionally KO (*Magel2* KO) as previously described^1^. Littermates expressing 2 wild type alleles were used as controls. Genotyping primers and protocols are as recommended by the manufacturer. For the *Magel2* WT allele: 5’-GTCACACACCCATTCGACCT-3’ and 5’-TACCCTCGGGAGCAGTAGAC-3’; *Magel2* KO allele: 5’-TGCTTCCTGCCCTTCAGTTAC-3’ and 5’-GCTTATCGATACCGTCGACCTC-3’; Cre-recombinase: 5’-TCTGTCCGTTTGCCGGTCGT-3’ and 5’-AGACCGCGCGCCTGAAGATA-3’; *Avp*-CRE WT allele: 5’-GAGTCCGTGGATTCTGCCAA-3’ and 5’-CTATGCACGACTTCGGGTGT-3’; *Oxt*-CRE WT allele: 5’-CTCAGAACACTGACCCATTTCTCTT-3’ and 5’-CCGACAATTAGACACCAGTCAAG-3’. All animals were housed under a 12/12 light/dark cycle in a SOPF sanitary status with access to ad libitum food and water. Enrichment consisted of cotton pieces to build a nest. Animals were group-caged until surgeries. After implantation surgery, animals were isolated until the end of the experiments. All efforts were made to minimize animal suffering and to reduce the number of mice utilized in each series of experiments.

### Behavior

The effects of *Magel2*^+m/−p^ mutation were evaluated on behaviors. A week before the 1^st^ experiment, mice (3-4 months old) were habituated to manipulation by the experimenter. A plexiglass transparent circular arena (24 cm diameter) was video-recorded for the time of the experiment and accommodated with clean litter, a water bottle and food before the beginning of the experiment. For social behavior, a transparent box with sniff holes was also introduced before the beginning of the experiment and a test mouse was left for 10 min to habituate to the arena. After 20 min of basal activity, a juvenile conspecific was introduced in the box for 5 min. The same presentation happened 4 times at 20 min intervals to habituate the mice with one another. After another 20 min, a new juvenile different from the first one was introduced in the box for 5 min. The set-up was systematically cleaned using disinfectant swipes before a new experimental mouse was challenged. This test in rodents typically decreases exploration of the known juvenile while it increases exploration of the new juvenile ^2^. For non-social behavior, a similar protocol was used except that an object was inserted into the arena. Time spent sniffing a juvenile freely moving or number of interactions with an object was extracted manually and expressed as % of interaction time during the first encounter.

#### *In vivo* pharmacology

Bilateral cannula (Phymed, 26G) heading to the lateral septum were implanted in mice using a stereotaxic frame (AP +0.05 cm, ML +/−0.04 cm, DV -0.3 cm) under ketamine (6.6 g/kg) xylazine (1.3 g/kg) anesthesia. Dental cement (Paladur, Henry Schein) and sutures were used to secure the implant. Animals were given anti-inflammatory medication (doliprane, 6 mg) and were monitored every day and allowed recovery for 7 days post-surgery. Positions of cannulas were verified *a posteriori* (**Fig. S9a**). Drugs were injected through the cannula using bilateral injector connected through 2 different tubing to two 1 μL Hamilton seringes controlled by a microinjector pumps (micro 4, World Precision Instrument). Injections of 900 μL/hemisphere were performed at 100 nL/min and diffusion perimeter was estimated by injection of Alexa-594-cadaverine (Life Technology, 50 μM, molecular weight comparable to the peptides used) using the same parameters just before euthanasia. The injection set-up was connected to the cannula 5 min before the injection and left for 5 min after the end of the injection such that it overlaps with the interaction with a mouse or object. The same mice were allowed at least 5 days between consecutive injections of NaCl 0.9 %, Atosiban (10^−8^ M in all experiments except 5.10^−8^ M for **fig 6a** and **6b** to disrupt OXTR during habituation as 2 consecutive injections at T1 and T3 impaired the robustness of the test), Manning compound (10^−8^ M), AVP (3×10^−6^ M) and TGOT (3×10^−7^ M) performed in random orders ^3,4^. Concentrations of aforementioned selective ligands were determined based on published selectivity profiles on mouse receptors ^5^. In brief, Atosiban is >700 times more potent on mOXTR than mAVPRs; TGOT is >20,000 times more potent on OXTR than mAVPRs; MC has a 24-42 times higher affinity for mAVPR1A than for mOXTR and mAVPR1B, respectively. Though AVP binds equally to mOXTR and mAVPR *in vitro* ^5^, it failed to elicit an electrophysiological response on OXT-excited type II neurons of brain slices. Thus, the priming effect of AVP on OXT response of inhibited-type III neurons is unlikely due to desensitization of OXTR.

### Optogenetics

*Oxt*-CRE, *Avp*-CRE, *Oxt*-CRE;*Magel2*^+m/−p^ (KO) and *Avp*-CRE;*Magel2*^+m/−p^ (KO) animals were used in these experiments. At 4 weeks, animal were injected with one of the following AAV1 (EF1a::DIO-eNpHR3.0-eYFP;WPRE::hGH; EF1a::DIO-ChR2-eYFP;WPRE::hGH or EF1a::DIO-eYFP;WPRE::hGH from viral vector core, University of Pennsylvania) (500 nL/hemisphere, 2×10^11^ viruses/mL) either in the hypothalamic paraventricular nucleus (AP -0.01 cm, ML +/−0.02 cm, DV 0.48 cm) or in the bed nucleus of the stria terminalis (BNST: AP 0.05 cm, ML +/−0.1 cm, DV -0.4 cm) using a microinjector-controlled (micro-4, WPI) seringe (nanofil seringe and Nanofil 33G BVLD Needle, WPI). One month after the infection, bilateral optic fibers (dual fiberoptic cannula, Doric lenses, 0.53 NA) were implanted in the septum (AP +0.05 cm, ML +/−0.04 cm, DV -0.3 cm). All surgeries were performed under ketamine (6.6 g/kg) xylazine (1.3 g/kg) anesthesia. LED were calibrated at 500 mA to deliver 2 mW at the tip of the optic fiber, as monitored with a luminometer (Thorlab). Stimulations consisted of 5 min continuous stimulation at 561 nm for the halorhodopsin and 5 ms pulses, 20 Hz or 10 ms pulses, 30 Hz at 473 nm for the channelrhodopsin (respectively for vasopressinergic or oxytocinergic neurons stimulation) as previously used ^6,7^. Positions of optic fibers were verified *a posteriori* (**Fig. S9b,c**).

### EEG recordings

Mice at least 2 months of age were implanted with surface electrodes atop the cortex and allowed two weeks of recovery from surgery before use for experiments. Electroencephalogram (EEG) activity was monitored for 3 h during interactions with a mouse or an object (Pinnacle Inc., Sarasota, FL, USA). EEG signals were acquired at 600Hz, stored and analyzed using Sirenia (Pinnacle Inc., Sarasota, FL, USA). Dual Video-EEG inspection permitted the exclusion of artifacts associated with scratching, eating, drinking, chewing, self-grooming, and sleep from interesting period of behavioral interactions with a conspecific or an object. Frequencies above 100Hz were filtered and data analyzed with Neuroscore software (Data Science International). Frequency bands of 0.5Hz were defined and power of each of these bands was evaluated for each 10s epoch composing the signal. Mean values of power for each band of frequency were calculated for the periods relevant with the behavior (while the social stimulus or the object is present in the arena). To limit the impact of inter-individual variability, data were normalized to a baseline measured for each animal just before the experiment started. Percent change of EEG power corresponds to the power spectrum values for each band of frequencies normalized to the area under the curve of the power spectrum considering frequencies between 1-100Hz (except the 48-52 Hz noisy band). Net change of EEG power in the theta band was calculated as follow: (power spectrum in 8-12 Hz band at T1, T4 or T5 - power spectrum in 8-12 Hz band at baseline) - (power spectrum in 4-8 Hz band at T1, T4 or T5 - % power spectrum in 4-8 Hz band at baseline).

### Live slice preparation

Animals (4-8 weeks old males) were anesthetized using isoflurane and quickly decapitated. Thick coronal slices (300 μm) containing the lateral septum were prepared using a vibrating microtome (Campden Instr. Ldt; England) and collected in an ice-cold slicing solution (in mM: 10 NaCl, 1.2 KCl, 26 NaHCO3, 15 glucose, 1.2 KH_2_PO_4_, 1 CaCl_2_, 2 MgCl_2_, 195 sucrose; osmolality adjusted to 300 mOsmol.l^−1^; pH 7.4, 95% O_2_ and 5% CO_2)_. Slices were allowed to recover for 1 h at 37°C in ACSF medium (in mM: 110 NaCl, 3.6 KCl, 26 NaHCO_3_, 10 glucose, 1.2 KH_2_PO_4_, 2 CaCl_2_, 2 MgCl_2_, 0.2 ascorbic acid, 0.2 thio-urea, osmolality adjusted to 300 mOsmol.l^−1^; pH 7.4, 95% O_2_ and 5% CO_2_) until placed in the recording chamber perfused at 1mL/min with heated ACSF.

### Slice electrophysiology

An Ag/AgCl electrode placed in the recording chamber was used as reference. A similar electrode was inserted in a borosillicated glass (GC150F-10, Harvard apparatus) electrode (4-6 Ohm resistance when immersed the bath) containing intracellular medium (in mM: 9 KCl, 130 KMeSO_3_, 8 NaCl, 1 MgCl_2_, 0.1 EGTA/Na, 10 HEPES/NaOH, 2 pyruvate, 2 malate, 0.5 NaH_2_PO_4_, 0.5 cAMP, 2 ATP-Mg, 0.5 GTP-Tris, 14 phosphocreatine, 0.1 leupeptine) and connected to the pre-amplifier. Recordings were performed using an axopatch 200B amplifier (Axon instruments; USA). Neurons visualized using infra-red Normarsky contrast microscopy and recorded from the lateral septum were chosen randomly. Bath application of peptides (TGOT 10^−7^ M or vasopressin 10^−6^ M) was performed in random order for 2 min to determine their impact on electrical/synaptic activity. GABAzine (10^−6^ M) or tetrodotoxine (TTX, 3×10^−7^M) were bath-applied for 10 min at least. Action potentials were recorded in whole-cell or cell-attached configurations for at least 5 min before any drug application, respectively in current or voltage clamp conditions. Interspike intervals (ISI) were calculated over a 3 min period and classified depending on their duration. Data were normalized to the duration of ISI. Alexa-Fluor-594-cadaverin (Life technology, 50 μM) was added to the intracellular medium of the patch pipette and whole-cell recordings performed as usual to determine cell morphology *a posteriori*.

### Retrograde tracings

Alexa-Fluor-555-conjugated latex microspheres (Retrobeads, Lumafluor) diluted 1:1 in PBS were stereotaxically injected in the medial septum (angle 10°, AP +0.05 cm, ML +0.1 cm, DV - 0.4 cm). After surgeries, mice were group-housed for at least a week before being sacrificed. Brains were processed for slice electrophysiology or immunohistochemistry.

### Immunohistochemistry

Mice were anesthetized with pentobarbital (50 mg/kg, i.p., Ceva Santé Animale, Libourne, France) and perfused at a rate of 3 ml/min through the ascending aorta with 30 ml of ice-cold 0.9% NaCl and 4 % ice-cold paraformaldehyde. Brains were sectioned with a vibratome and free-floating coronal sections rinsed in PBS were blocked in 3 % normal donkey serum, PBS, 0.1 % triton X-100 for 2 h at 25 °C. Non-labelled Fab anti mouse IgG (abliance, 1:500) was used to block non-specific sites when mouse antibodies were chosen. Primary antibodies (dilutions for: NG 1:1000, Synaptophysin 1:100, c-Fos 1:1000, p-S6 1:400, Som 1:50, Neurotensin 1:250, GAD67 1:500, NeuN 1:500, Calbindin 1:5000, Calretinin 1:1000, Neurphysin I 1:1000, Neurophysin II 1:500) were incubated overnight. Alexa-Fluor-conjugated secondary antibodies (1:2,000 ThermoFisher Scientific, Waltham, MA, USA) were incubated for 2 h at 25°C. Images were acquired with a confocal microscope LSM780 (Carl Zeiss, Iena, Germany) and 10x, 20x dry objectives or 40x oil-immersion objective for co-expression studies and with a epifluorescence microscope (Imager.Z1, axiocam, Carl Zeiss, Iena, Germany). Excitation and acquisition parameters were unchanged during the acquisition of all images.

### Image analyses

More than 67500 cFos cells, 6727 OXT neurons, 5978 AVP neurons, 1665 RB-555 containing cells, and 4491 d[Lys(Alexa-Fluor-647)^8^]VP labeled cells were counted in all groups to determine proportions of various cells types co-labeled with c-Fos. Images spanning the anterior, central and posterior septum from the same mouse served to count the number of c-Fos positive cells normalized to the surface of the region of interests marked with Image J software (NIH). C-Fos+ cells were further averaged between groups of mice for comparison. Reconstruction of the dendritic arborization in Cadaverine-filled cells from Z-stack images (1μm steps) and Scholl analysis was performed using Image J. Semi-guided tracing of OXT and AVP fibers into the septum was performed with the NeuronJ plugin. Positions of cannulas and fibers optics were determined under the microscope in at least 3 sections of septum in each postmortem brain. The diffusion areas of the Alexa-594-cadaverine dye were compared in fixed sections of septum between animals to determine *a posteriori* the diffusion area of drugs with similar molecular weight (*e.g.* AVP, TGOT, MC, Atosiban).

### Synthesis of d[Lys(Alexa-Fluor-647)^8^]VP

The synthesis of the fluorescent peptide was carried out in the laboratory of Dr M. Manning (University of Toledo, OH) as previously described ^8^. All solvents and reagents used were analytical grade. Most standard chemicals were purchased from Sigma (St. Louis, MO) or Merck (Darmstadt, Germany). [Deamino-Cys^1^, Lys^8^] Vasopressin (Bachem, Bubendorf, Switzerland) 1 mg in 100 μL of anhydrous dimethyl formamide and 2.5 μL (14 μmol) of DIPEA (*N,N*-diisopropylethylamine) was added to 1 mg (1.25 *μ*mol, MW ~ 800) of the Alexa-Fluor®-647 carboxylic acid succinimidyl ester (Invitrogen/ThermoFisher Scientific). The reaction mixture was stirred for 5 h at room temperature in the dark, then mixture was acidified (6 μL TFA), diluted with 10 ml of water and lyophilized. The crude material was diluted with 3 mL of 0.05% TFA in water, purified by semi-preparative HPLC on a Hitachi D-7000 HPLC system (detection at 214 nm), using a C18 column (218TP510 C18, Vydac hromatograph Grace Vydac, Hesperia, CA), solvent A 0.05% Tri-Fluoro Acetic acid (TFA) in water, solvent B 0.05%TFA in acetonitrile, gradient 10-70%B over 30 min, flow rate 5.0 mL/min. Yield of lyophilized dLVP 1.5 mg (~87%), RP-HPLC: t_R_15.32 min, purity 100%. The affinity of d [Lys (Alexa-Fluor-647)^8^] VP was tested by competitive binding assays with [H]^3^AVP on membrane preparations of HEK293 cells transfected with recombinant mOXTR or mAVPR1B, and liver or kidney to purify endogenous mAVPR1A and AVPR2 as previously described ^8^. All assays were run in triplicates in at least n=3 independent experiments. The selectivity profile of d[Lys(Alexa-Fluor-647)^8^]VP is best for mOXTR as its Ki affinity was higher 54, 205 and 2796 times than that of AVPR1B, mAVPR1A and mAVPR2, respectively. A similar strategy was previously used to characterize V1b expressing cells in the brain of rats ^9^.

### Statistics

Parameters quantified include (i) exploration time with a mouse or an object, (ii) frequency of spikes or synaptic events, (iii) co-expression of d[Lys(Alexa-Fluor-647)^8^]VP with cell type markers, (iv) position of cannulas, optic fibers and viral-mediated recombination at injection sites, (v) density of c-Fos cells in LS, (vi) change of EEG activity between epochs. Representation of n for each data set is indicated in figure legends. All data collected in animals were from littermate controls and averaged per experimental groups. Data were considered non-parametric when sample size was moderate. Data were tested for normality and compared to a theorical value with Wilcoxon signed rank test (Prism 8.0 Software, GraphPad). We used a paired t-test for comparing 2 series of values from the same animal or cell. Otherwise we used Mann Whitney t-test. We used Spearman correlation for linear associations between datasets. Data considering one variable and more than 2 conditions were analysed using 1-way ANOVA (Kruskal Wallis). Dunns post-hoc test were used to compare the different groups. We used factorial ANOVA to compare multiple groups (behavior, genotype, optogenetics and pharmacology) followed by *post-hoc* pairwise comparison with Sidak, Dunnett or Tukey tests for corrections. All data are shown as means ± standard error of the mean. Significance level is set at α≤0.05. Randomization: for in vivo pharmacology experiments, animals received the different compounds in a random order. For slice electrophysiology experiments, TGOT or vasopressin were administered in a random order. Inclusion or exclusion criteria: (i) postmortem validation of positions of cannulas, (ii) optic fibers and (iii) viral-mediated recombination. An independent experimenter blind to the conditions performed these analyses. Pre-established criteria for excluding data: (i) misposition of cannulas or fibers optics; (ii) poor recombination of viral-guided tools; (iii) instability of EEG signal. Estimates of sample size were calculated by power analysis based on preliminary data. Sample size was chosen to ensure 80 % power to detect the pre-specified effect size. Pre-established criteria for stopping data collection included: (i) mice reaching ethical endpoint limits; (ii) unexpected detachment of cranial implants; (iii) mice interacting < 15 s during the 1^st^ social encounter or < 10 times during the 1^st^ encounter with the object; (iv) brains badly perfused and unusable for histology. Complete statistic information can be found in **Supplemental Table 1**.

## Data availability

Data and reagents can be shared on demand.

## Supplemental Figures

**Figure S1.**
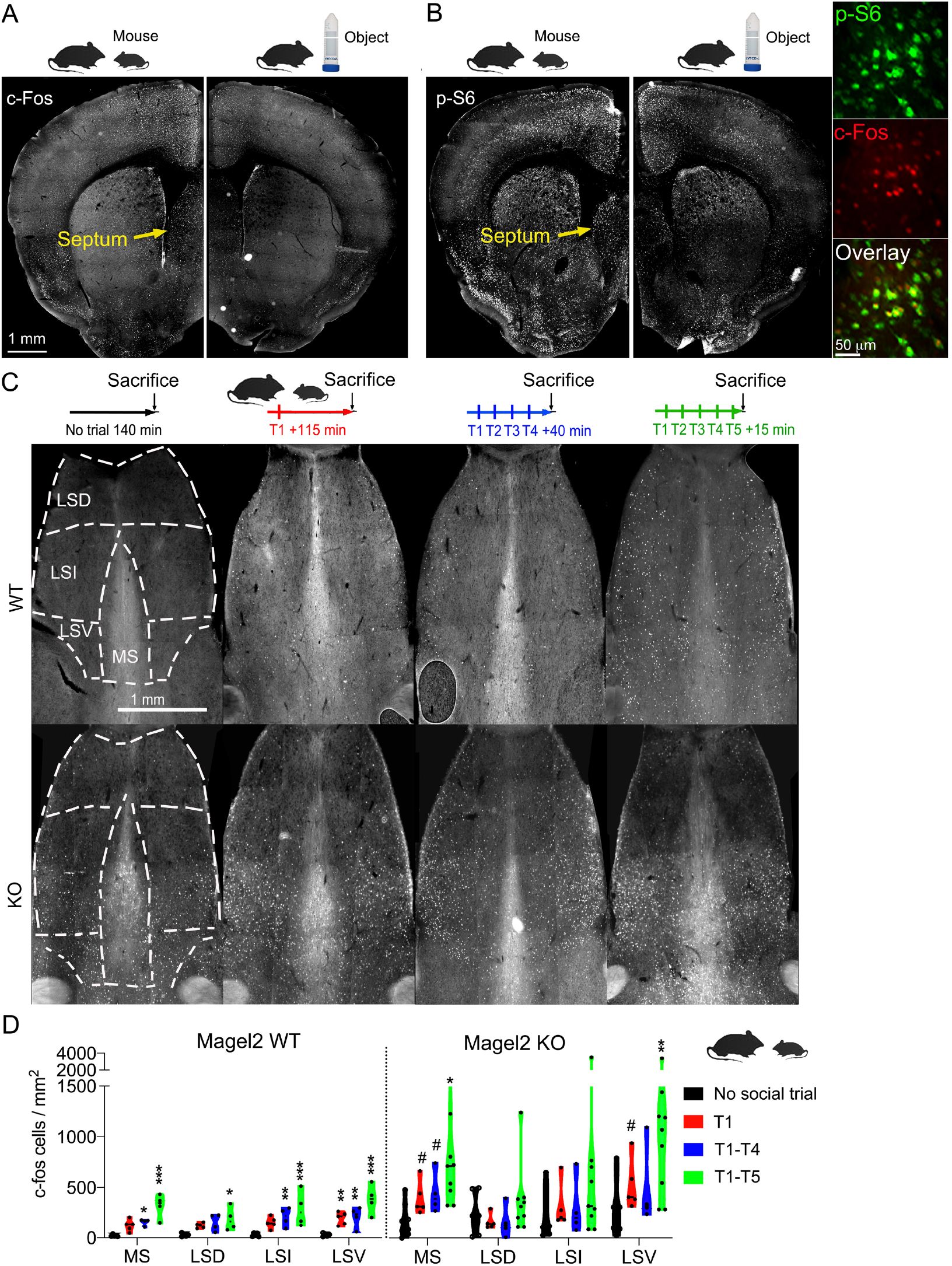
Fos mapping of septum reactivity to social trials of *Magel2^+m/−p^* mice with new WT conspecifics. (**a**) Representative c-Fos immunoreactivity in a WT mouse interacting with a new WT conspecific for 5 min, sacrificed 15 min later (left) compared to a WT mouse interacting with a new object in the same conditions (right). (**b**) Representative p-S6 immunoreactivity in a WT mouse interacting with a new WT conspecific for 5 min, sacrificed 15 min later (left) compared to a WT mouse interacting with a new object in the same conditions (right). Colored insets represent a zoom in the dorsal lateral septum co-labeled with c-Fos and p-S6. Same cells responded with both markers in mice sacrificed 15 min after social trial except that signal detection was more robust with an established marker of rapid signaling (p-S6) than a c-Fos, a surrogate of immediate early gene transcription ^10^. (**c**) Close up look at septal c-Fos immunoreactivity to 1 or more social trials with new conspecifics in *Magel2^+m/−p^* and WT mice as described in Figure 1A. Experimental timelines are as follow: T0 = 140 min in test chamber before sacrifice; T1 = 1 interaction followed by 115 min in test chamber before sacrifice; T1-T4 = 4 interactions followed by 40 min in test chamber before sacrifice; T1-T5 = 5 interactions followed by 15 min in test chamber before sacrifice. (**d**) Density of c-Fos positive cells/mm^2^ in a total of 28072 WT and 38534 KO cells. Means ± SEM of n = 6 T0, 5 T1, 4 T4, 4 T5 WT mice and 7 T0, 5 T1, 4 T4, 9 T5 *Magel2^+m/−p^* KO mice. Two-way ANOVA: Effect of social trials in WT: *F*(3, 56) = 31.7, *p* < 0.0001 post-hoc Dunnett test in MS: T0 vs T5 \*\*\**p* < 0.0001 and T0 vs T4 \**p* = 0.045; in LSD: T0 vs T5 \**p* = 0.016; in LSI: T0 vs T4 \*\**p* = 0.0063, T0 vs T5 \*\*\**p* < 0.0001; in LSV: T0 vs T1 \*\**p* = 0.007, T0 vs T4 \*\**p* = 0.0058, T0 vs T5 \*\*\**p* < 0.0001. Two-way ANOVA: Effect of social trials in KO: *F*(3, 88) = 6.871, *p* = 0.0003 post-hoc Dunnett test in MS: T0 vs T5 \**p* = 0.0357; in LSV: T0 vs T5 \*\**p* = 0.0025. Multiple comparison between WT and KO by t-test at T1 in MS *t*(8) = 3.5 ^#^*p* = 0.007 and LSV *t*(8) = 2.8 ^#^*p* = 0.02; at T4 in MS *t*(6) = 2.76 ^#^*p* = 0.032.

**Figure S2.**
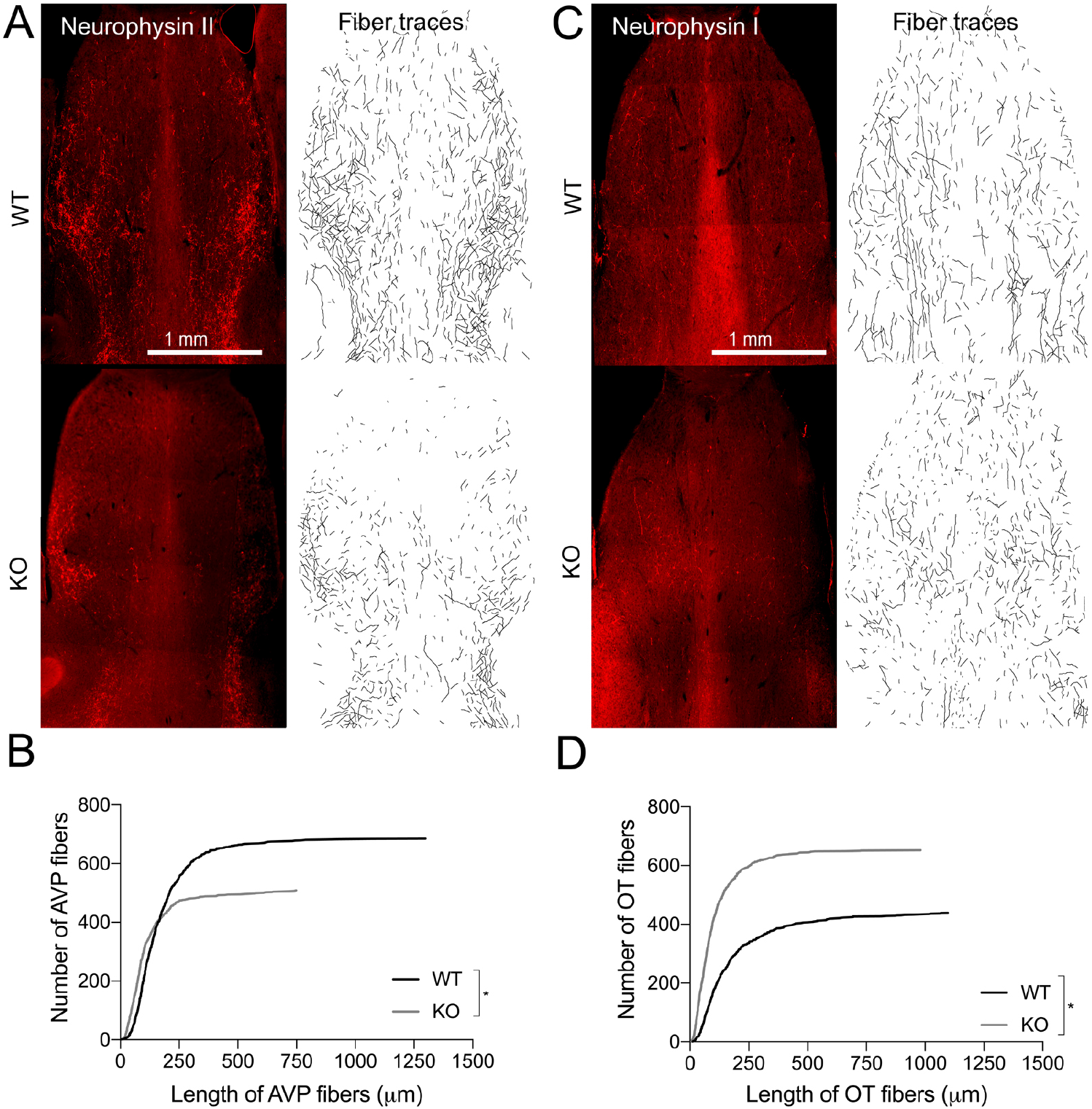
Innervation of AVP and OXT fibers in septum of WT and *Magel2^+m/−p^* KO mice. (**a**) Representative immunoreactivity of neurophysin II (AVP) in septum of WT and *Magel2^+m/−p^* mice. Semi-automated tracing of AVP fibers are shown using NeuronJ. (**b**) Number of AVP fibers in 5 WT and 5 KO mice sorted as a function of the length. Kolmogorov-Smirnov test *p* < 0.0001. (**c**) Representative immunoreactivity of neurophysin I (OXT) in septum of WT and *Magel2^+m/−p^* mice. Semi-automated tracing of OXT fibers are shown using NeuronJ. (**d**) Number of OXT fibers in 5 WT and 5 KO mice sorted as a function of the length. Kolmogorov-Smirnov test *p* < 0.0001.

**Figure S3.**
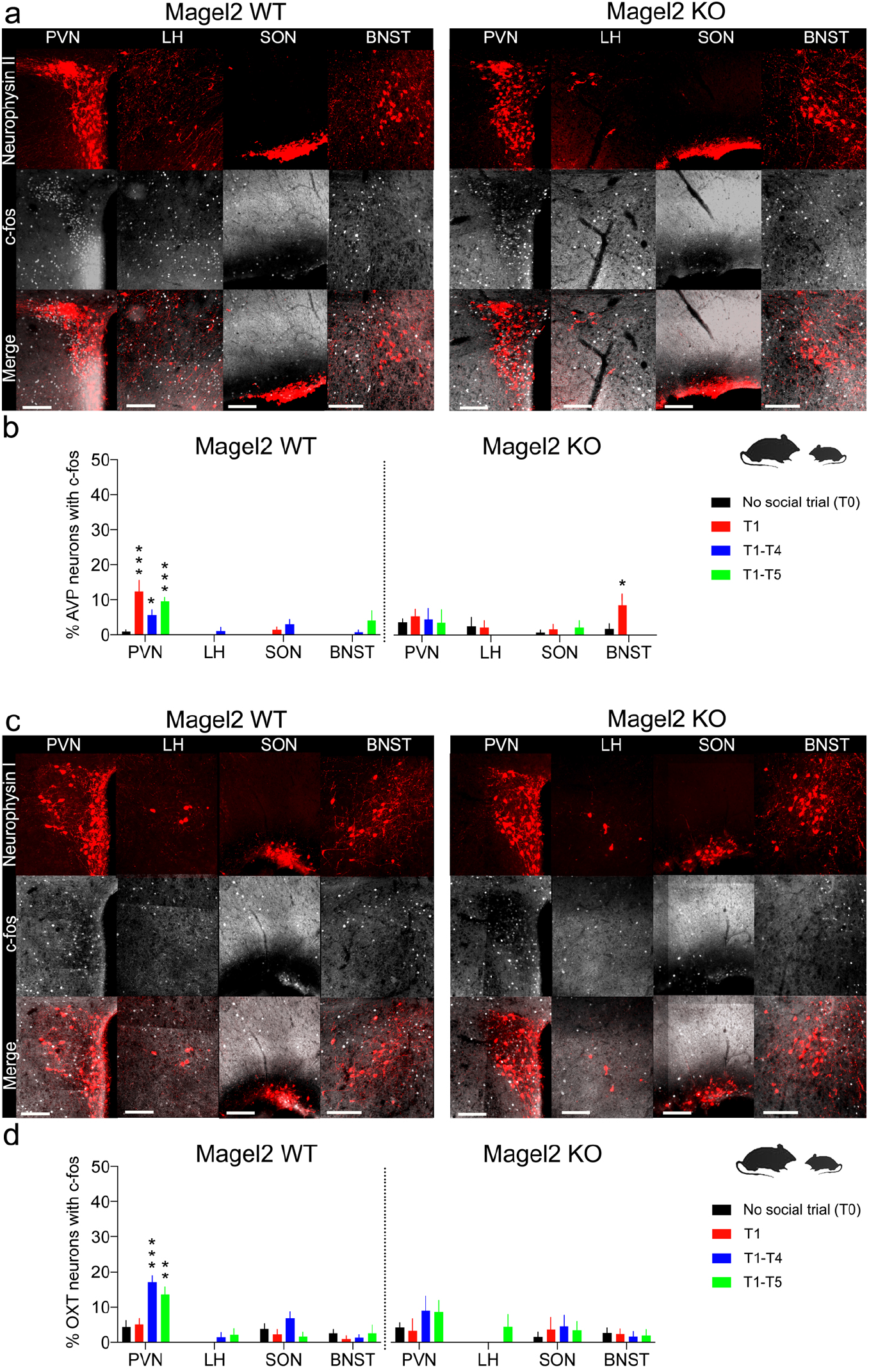
Fos mapping of OXT and AVP neuron reactivity to social trials with new WT conspecifics in *Magel2* WT and KO mice. (**a**) Induction of c-Fos in AVP neurons (neurophysin II) of paraventricular hypothalamic nucleus (PVN), lateral hypothalamus (LH), supraoptic nucleus (SON) and bed nucleus of stria terminalis (BNST) after 1 social trial (T1) in *Magel2^+m/−p^* mice and WT controls. Experimental timelines are as follow: T0 = 140 min in test chamber before sacrifice; T1 = 1 interaction followed by 115 min in test chamber before sacrifice; T1-T4 = 4 interactions followed by 40 min in test chamber before sacrifice; T1-T5 = 5 interactions followed by 15 min in test chamber before sacrifice. Scale bars = 50 μm. (**b**) Percentage of 4133 WT, 1845 KO AVP neurons expressing c-fos as a function of social trials. Means ± SEM of n = 5 T0, 5 T1, 4 T4, 4 T5 WT and *Magel2^+m/−p^* mice. Two-way ANOVA: Effect of social trials on regional AVP neurons in WT: *F*(3, 56) = 24.61, *p* < 0.0001 post-hoc Dunnett test in PVN: T0 vs T1 \*\*\**p* < 0.0001, T0 vs T5 \*\*\**p* < 0.0001 and T0 vs T4 \**p* = 0.02; in KO: *F*(3, 56) = 2.15, *p* = 0.1 post-hoc Dunnett test in BNST: T0 vs T1 \**p* = 0.038. (**c**) Induction of c-Fos in OXT neurons (neurophysin I) of PVN, LH, SON and BNST after 4 social trial (T1-T4) in *Magel2^+m/−p^* mice and WT controls. Scale bars = 50 μm. (**d**) Percentage of 4730 WT, 1997 KO OXT neurons expressing c-fos as a function of social trials. Means ± SEM of n = 5 T0, 5 T1, 4 T4, 4 T5 WT and *Magel2^+m/−p^* mice. Two-way ANOVA: Effect of social trials on regional AVP neurons in WT: *F*(3, 56) = 29.82, *p* < 0.0001 post-hoc Dunnett test in PVN: T0 vs T4 \*\*\**p* < 0.0001, T0 vs T5 \*\**p* = 0.002; in KO: *F*(3, 56) = 3.84, *p* = 0.0143.

**Figure S4.**
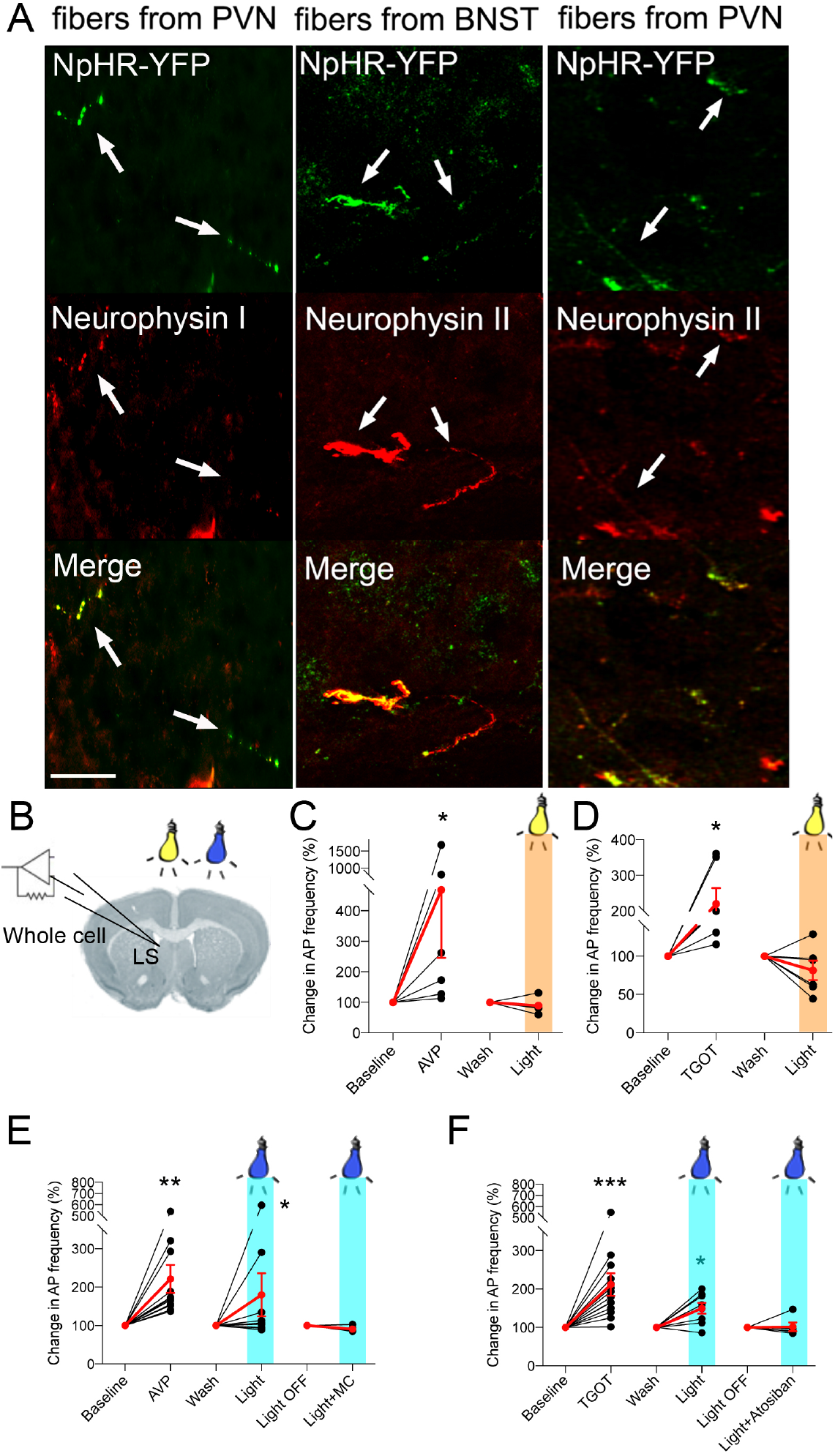
Optogenetic control of AVP and OXT receptor responses in LS. (**a**) Viral-mediated expression of NpHR3.0-YFP in PVN to LS fibers of OXT neurons in *Oxt*-CRE as well as in PVN to LS fibers or BNST to LS fibers of AVP neurons in *Avp*-CRE mice. Scale = 25 μm. (**b**) Spontaneous activity of LS cells recorded in whole cell configuration in brain slices of recombinant mice. (**c**) Effect of yellow light stimulation (561 nm, 5 ms pulses, 20 Hz for 2 min, ~2 mW) on the relative frequency of action potentials (AP) of cells pre-identified as AVP-excited (10^−6^ M, 2 min). n = 7 cells (Means ± SEM in red): Wilcoxon test \**p* = 0.015. (**d**) Effect of yellow light stimulation (561 nm, 5 ms pulses, 20 Hz for 2 min, ~2 mW) on the relative AP frequency of cells pre-identified as TGOT-excited (10^−8^ M, 2 min). n = 6 cells (Means ± SEM in red): Wilcoxon test \**p* = 0.031. (**e**) Effect of blue light stimulation (473 nm, 20 Hz, 5 ms pulses for 2 min, ~2 mW) on the relative AP frequency of cells pre-identified as AVP-excited (10^−6^ M, 2 min). n = 7 cells without MC and 5 cells with MC (10^−8^ M) (Means ± SEM in red): Wilcoxon test \*\**p* = 0.001 and \**p* = 0.046. (**f**) Effect of blue light stimulation (473 nm, 30 Hz, 10 ms pulses for 2 min, ~2 mW) on the relative AP frequency of cells pre-identified as TGOT-excited (10^−7^ M, 2 min). n = 8 cells without atosiban and 5 cells with atosiban (10^−8^ M) (Means ± SEM in red): Wilcoxon test \*\*\**p* < 0.0001 and \**p* = 0.02.

**Figure S5.**
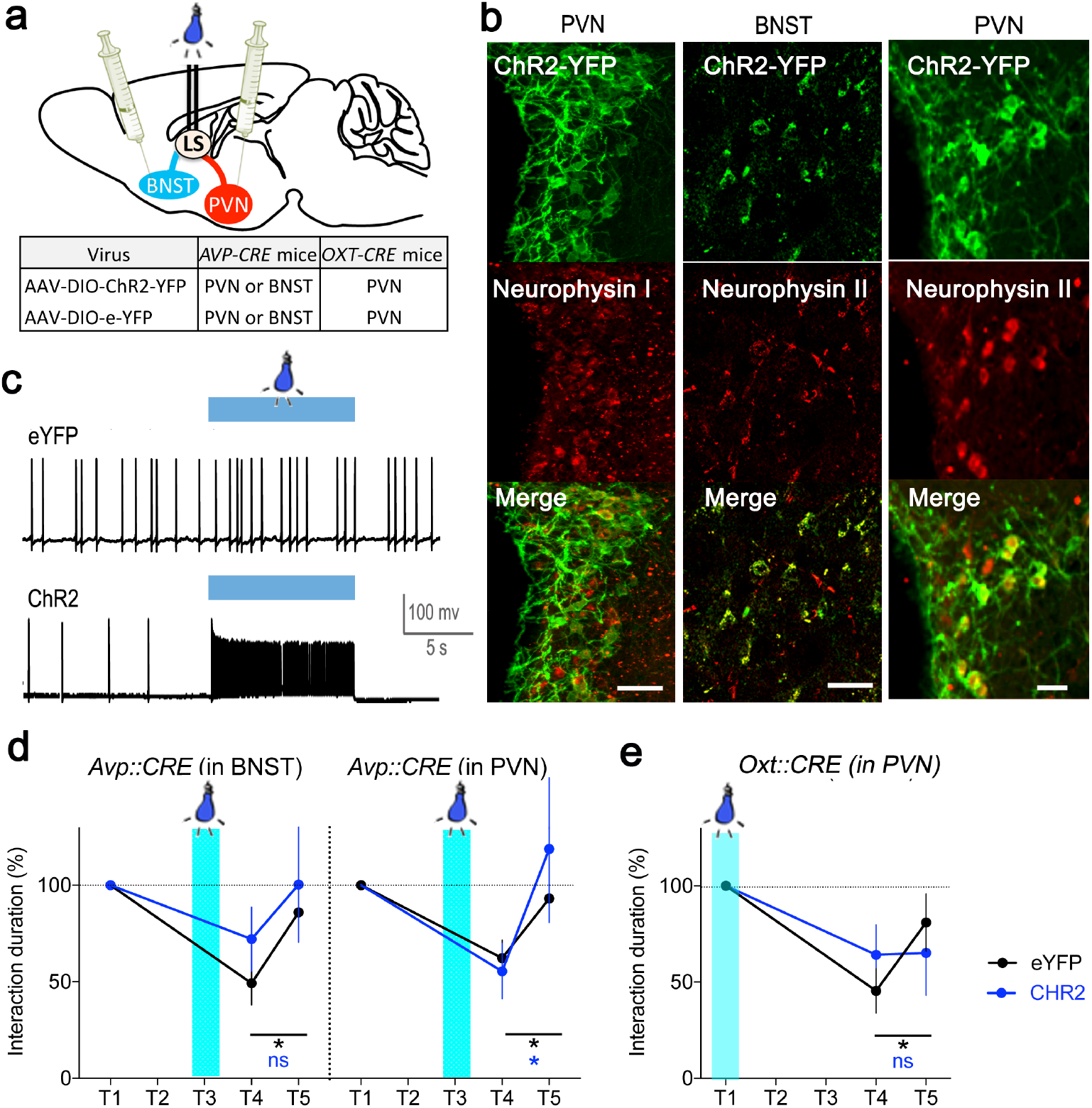
Optogenetic-mediated disorder of AVP and OXT sequential responses in LS impaired social discrimination. (**a**) Viral-mediated optogenetic stimulation of septal inputs from OXT neurons (*Oxt*-CRE mice) in PVN or from AVP neurons (*Avp*-CRE mice) in PVN or BNST. (**b**) Co-expression of Cre-mediated ChR2-YFP with Neurophysin I (OXT) in PVN neurons or with Neurophysin II (AVP) in PVN as well as BNST neurons. Scale = 40 μm. (**c**) Firing of action potentials recorded in whole cell configuration in neurons expressing ChR2-YFP or its eYFP. Stimulation with blue light (473 nm, 30 Hz, 10 ms pulses for 5 min, ~2 mW) increased firing rate of ChR2 cells. (**d**) Percent time interacting with a mouse throughout trials. Data (means ± SEM) are relative to T1 in each group of n = 12 eYFP, 8 ChR2 in BNST and 9 eYFP, 6 ChR2 in PVN of *Avp-CRE* mice. Two-way ANOVA: Effect of ChR2 stimulation of BNST fibers by blue light (473 nm, 20 Hz, 5 ms pulses for 2 min, ~2 mW at T3): *F*(2, 36) = 6.76, *p* = 0.0032 post-hoc Dunnett test for eYFP controls comparing T4 with T1 \*\**p* = 0.0024, and T4 with T5 \**p* = 0.028; Effect of ChR2 stimulation of PVN fibers by blue light (473 nm, 20 Hz, 5 ms pulses for 2 min, ~2 mW at T3): *F*(1, 13) = 0.2, *p* = 0.65 post-hoc Dunnett test for eYFP controls comparing T4 with T1 \*\**p* = 0.007, and T4 with T5 \**p* = 0.022; and for ChR2 comparing T4 with T1 \**p* = 0.046, and T4 with T5 *p* = 0.18. (**e**) Percent time interacting with a mouse throughout trials. Data (means ± SEM) are relative to T1 in each group of n = 12 eYFP, 7 ChR2 in PVN of *Oxt-CRE* mice stimulated with blue light in LS (473 nm, 30 Hz, 10 ms pulses for 2 min, ~2 mW at T3). Two-way ANOVA: Effect of social trials: *F*(2, 34) = 6.77, *p* = 0.0033, post-hoc Dunnett test for eYFP controls comparing T4 with T1 \*\**p* = 0.0018, and T4 with T5 \**p* = 0.043.

**Figure S6.**
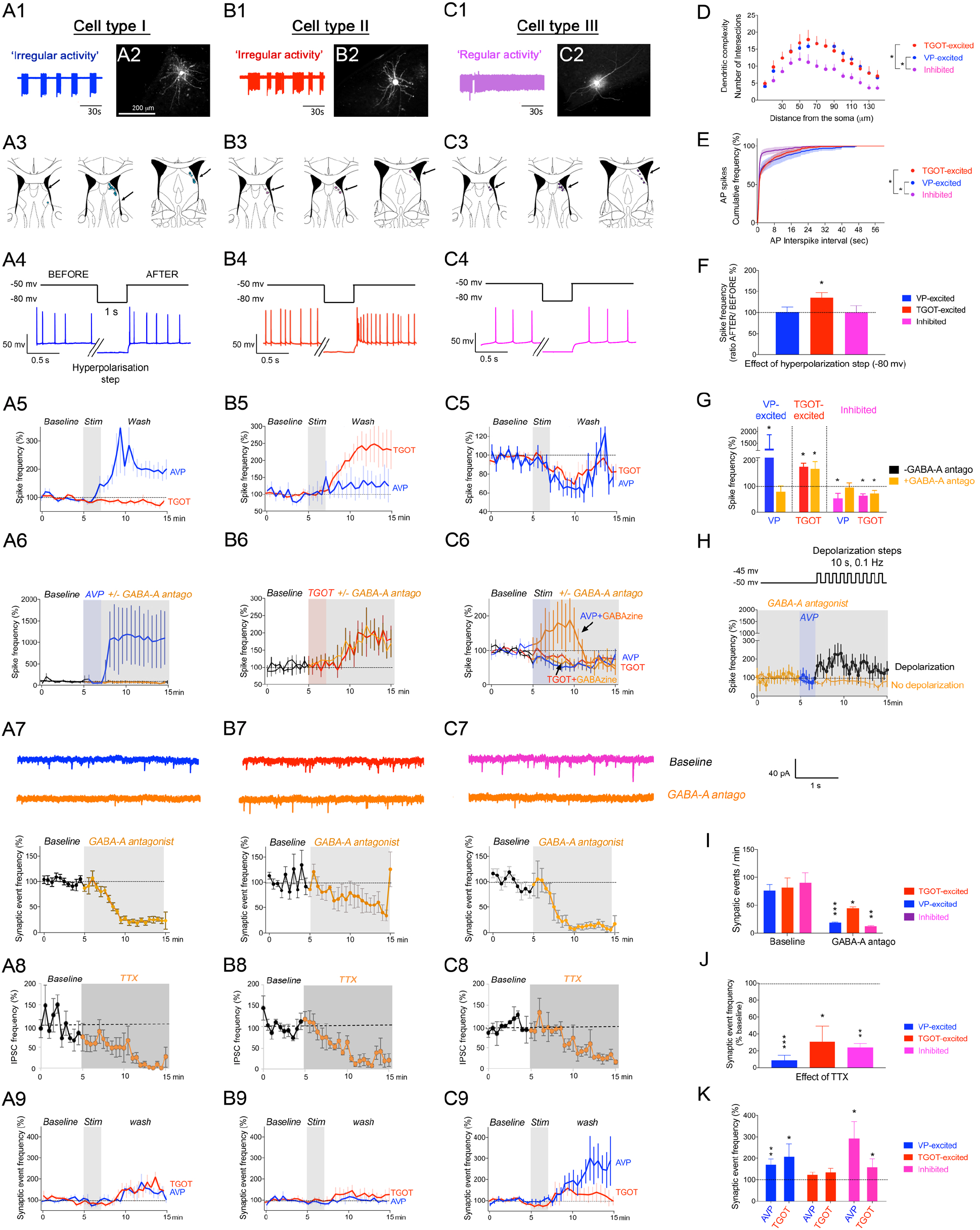
Characterization of 3 major populations of neurons in LS based on their electrophysiological responses to AVPR and OXTR agonists. *In brief, type I cells are bursting neurons under strong GABA-A dependent AVP regulation; type II are irregular bursting neurons with a hyperpolarization-activated current whose excitation by TGOT is GABA-A independent; and type III neurons exhibit sustained activity decreased by AVP (GABA-A dependent) or by TGOT (GABA-A independent).* (**a**) Type I: spontaneous spike activity in bursts (A1) of branched cells marked with cadaverine-Alexa594 (A2, see D for group analysis) randomly distributed in LS (A3), excited only by 2 min of 10^−6^ M AVP (A5) and completely dependent on GABA-A receptors (A6, see G for group analysis). Synaptic inputs are mostly GABAergic (A7, see I for group analysis) coming from local cells (A8, see J for group analysis); their frequency is increased by TGOT and AVP (A9, see K for group analysis). (**b**) Type II: spontaneous spike activity irregular in bursts (B1) of branched cells (B2, see D for group analysis) randomly distributed in LS (B3), excited only by 2 min of 10^−7^ M TGOT (B5) and completely independent on GABA-A receptors (B6, see G for group analysis). Spike activity on these cells is increased by a hyperpolarization-activated current step (−80 mv, 1 s) (B5, see F for group analysis). Synaptic inputs both GABAergic and non-GABAergic (B7, see I for group analysis) most of which come from local cells (B8, see G for group analysis); their frequency is not sensitive to TGOT or AVP (B9, see K for group analysis). (**c**) Type III: Regular and sustained spike activity (C1) of tripolar cells (C2, see D for group analysis) randomly distributed in LS (C3), inhibited by AVP and also by TGOT (C5). Spike pattern on these cells is interrupted with silences introduced by AVP and TGOT (C4, see F for group analysis). AVP effect depends on GABA-A receptors contrary to TGOT effect that is insensitive (C6, see G for group analysis). Synaptic inputs are mostly GABAergic (C7, see I for group analysis) coming from local cells (C8, see J for group analysis); their frequency is increased by AVP only (C9, see K for group analysis). (**d**) Morphological characterization of each cell type patched and filled with 50 μM cadaverine-Alexa594 for Scholl analysis. Means ± SEM of n = 10 type I cells, 8 type II cells and 7 type III cells. One-way ANOVA: Effect of cell type: *F*(2, 39) = 6.91, *p* = 0.0027 post-hoc Tukey test: Type I vs type III \**p* = 0.0072 and Type II vs type III \**p* = 0.007. (**e**) Patterns of activity are distinct between cell types. Cumulative frequency of the time during which the cell fires at a frequency defined by each interspike interval epoch. Two-way ANOVA: Effect of cell type: *F*(2, 12080) = 91.19, *p* < 0.0001 post-hoc Tukey test: \**p* < 0.0001. (**f**) Bursting activity of type II cells modified by a hyperpolarization-activated current. Hyperpolarization step (−80 mv, 1 s) increased spike frequency immediately after repolarization only in type II cells. Means ± SEM of n = 20 type I, 12 type II and 11 type III cells. Paired t-test *t*(11) = 3.022, \**p* = 0.011. (**g**) Average spike frequency in each cell type. Means ± SEM of n = 7 type I, 6 type II and 7 type III cells. Wilcoxon paired t-test: Effect of 10^−6^ M AVP on type I cells (\**p* = 0.0156) lost with GABAzine; No effect of 6 10^−6^ M GABAzine on type II cells stimulated with 10^−7^ M TGOT (\**p* = 0.031); Effect of AVP on type III cells (\**p* = 0.046) and TGOT on type III cells (\**p* = 0.015) were lost with GABAzine. (**h**) Inhibitory effect of 6 10^−6^ M GABAzine on AVP-mediated excitation of type I cells relieved by small depolarization steps (+5mv, 10 s, 0.05 Hz). Therefore, AVP action depends on the existence of small depolarizing events whose impact on neuronal spiking activity is probably increased by AVP (A4). Means ± SEM of n = 9 type I cells with depolarization steps and 8 without. (**i**) Effect of 6 10^−6^ M GABAzine on the frequency of synaptic events. Means ± SEM of n = 14 type I, 8 type II and 6 type III cells. Two-way ANOVA: *F*(1, 50) = 44.8, *p* < 0.0001 post-hoc Sidak test on type I \*\*\**p* < 0.0001, type II \**p* = 0.05 and type III \*\**p* = 0.0002. (**j**) Effect of 3 10^−7^ M TTX on the frequency of synaptic events. Means ± SEM of n = 7 type I, 6 type II and 3 type III cells, paired t-test \*\*\**p* < 0.0001, \*\**p* = 0.0034, \**p* = 0.013. (**k**) Effect of TGOT and AVP on the frequency of synaptic events in each cell type. n = 15 type I, 9 type II and 10 type III cells. Paired t-test for type I *t*(9) = 3.36, \*\**p* = 0.0028 and *t*(9) = 2.25, \**p* = 0.04; type III *t*(8) = 2.52, \**p* = 0.035 and *t*(9) = 2.79, \**p* = 0.02.

**Figure S7.**
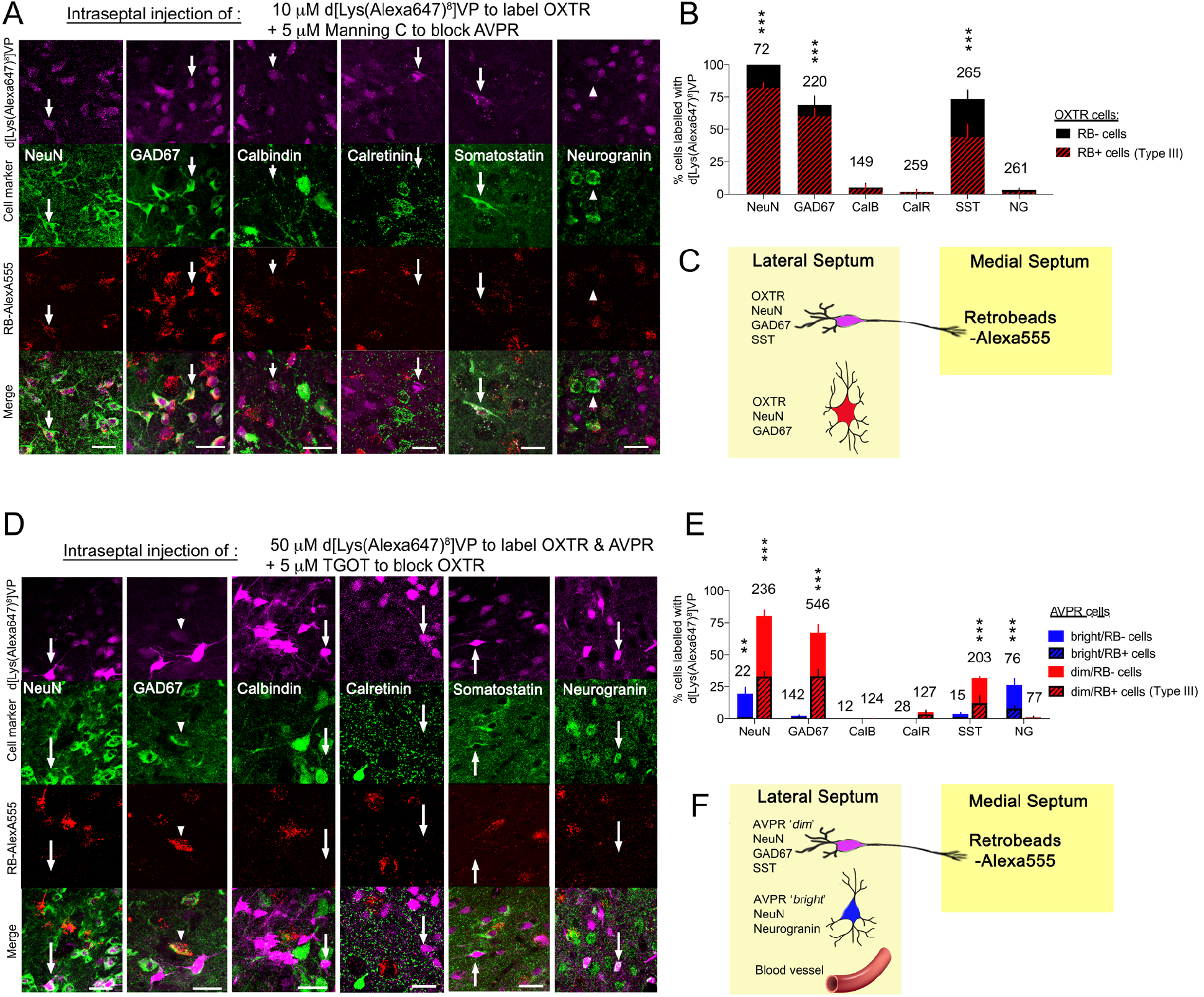
Type III cells are GABAergic neurons projecting to MS, equipped with AVPR and/ or OXTR. (**a**) Co-expression of OXTR binding sites [injection of d[Lys(Alexa-Fluor-647)^8^]VP +MC] with the indicated cellular markers. Arrows point to cells equipped with OXTR. Arrowheads point to cells not equipped with OXTR. Scale bars = 25 μm. (**b**) Percent of cells equipped with OXTR and the indicated marker. Number of cells in specific categories harboring OXTR is indicated out of 1226 OXTR cells of which 334 RB+. Wilcoxon unpaired t-test to evaluate the significance of co-expression of OXTR with other markers as well as OXTR/RB+ with other markers \*\*\**p* < 0.0001. (**c**) LS cells equipped with OXTR are GABAergic/ SST+ and can be sorted in 2 categories: with or without retrobeads. (**d**) Co-expression of AVPR binding sites [injection of d[Lys(Alexa-Fluor-647)^8^]VP +TGOT] with the indicated cellular markers. Arrows point to “bright” cells and arrowheads point to “dim” cells equipped with AVPR. Scale bars = 25 μm. (**e**) Percent of cells equipped with AVPR and the indicated marker out of 1608 AVPR cells of which 523 RB+. Number of cells in specific categories harboring AVPR is indicated. Wilcoxon unpaired t-test to evaluate the significance of co-expression of AVPR with other markers as well as AVPR/RB+ with other markers \*\*\**p* < 0.0001, \*\**p* < 0.001. (**f**) LS cells equipped with AVPR can be sorted in 4 categories: GABAergic neurons with or without retrobeads and SST, glutamatergic neurons without retrobeads, and blood vessels (see **Figure 5b**).

**Figure S8.**
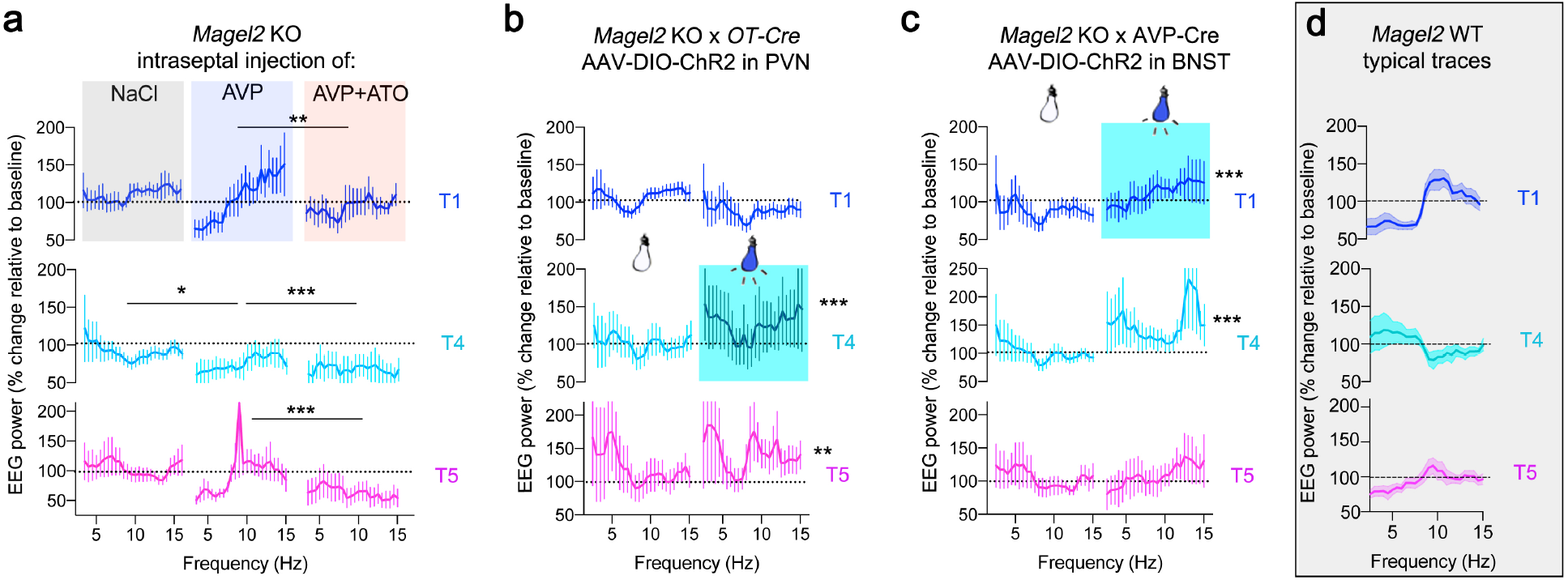
Optogenetic and pharmacological manipulations of *Magel2^+m/−p^* KO EEG activity during social trials. (**a**) Percent change of EEG power in 4-12 Hz band upon bilateral injection of 900 nL of NaCl, 3.10^−6^ M AVP with or without 5.10^−8^ M atosiban at T1 directly in septum of *Magel2^+m/−p^* mice. Means ± SEM of n = 13 NaCl, 12 AVP and 6 AVP+atosiban mice. Kruskal Wallis test at T1 *t*(3) = 16.47, *p* = 0.0003, effect of AVP vs AVP+ATO \*\**p* = 0.0063; at T4 *t*(3) = 41.61, *p* < 0.0001, effect of AVP vs NaCL \**p* = 0.04, AVP vs AVP+ATO \*\*\**p* < 0.0001; at T5 *t*(3) = 34.71, *p* < 0.0001, effect of AVP vs AVP+ATO \*\*\**p* < 0.0001. (**b**) Percent change of EEG power in 4-12 Hz band upon bilateral blue light stimulation (473 nm, 30 Hz, 10 ms pulses for 2 min, ~2 mW) in LS of OXT neuron projections between PVN and LS (PVN-LS pathway). Means ± SEM of n = 12 no stim, 12 stim mice. Mann Whitney test at T4 \*\*\**p* < 0.0001, at T5 \*\**p* = 0.0031. (**c**) Percent change of EEG power in 4-12 Hz band upon bilateral blue light stimulation (473 nm, 30 Hz, 10 ms pulses for 2 min, ~2 mW) in LS of AVP neuron projections between BNST and LS (BNST-LS pathway). Means ± SEM of n = 9 no stim, 6 stim mice. Mann Whitney test at T1 and T4 \*\*\**p* < 0.0001. (**d**) Typical theta rhythm activity traces socially evoked in control WT mice for comparisons.

**Figure S9.**
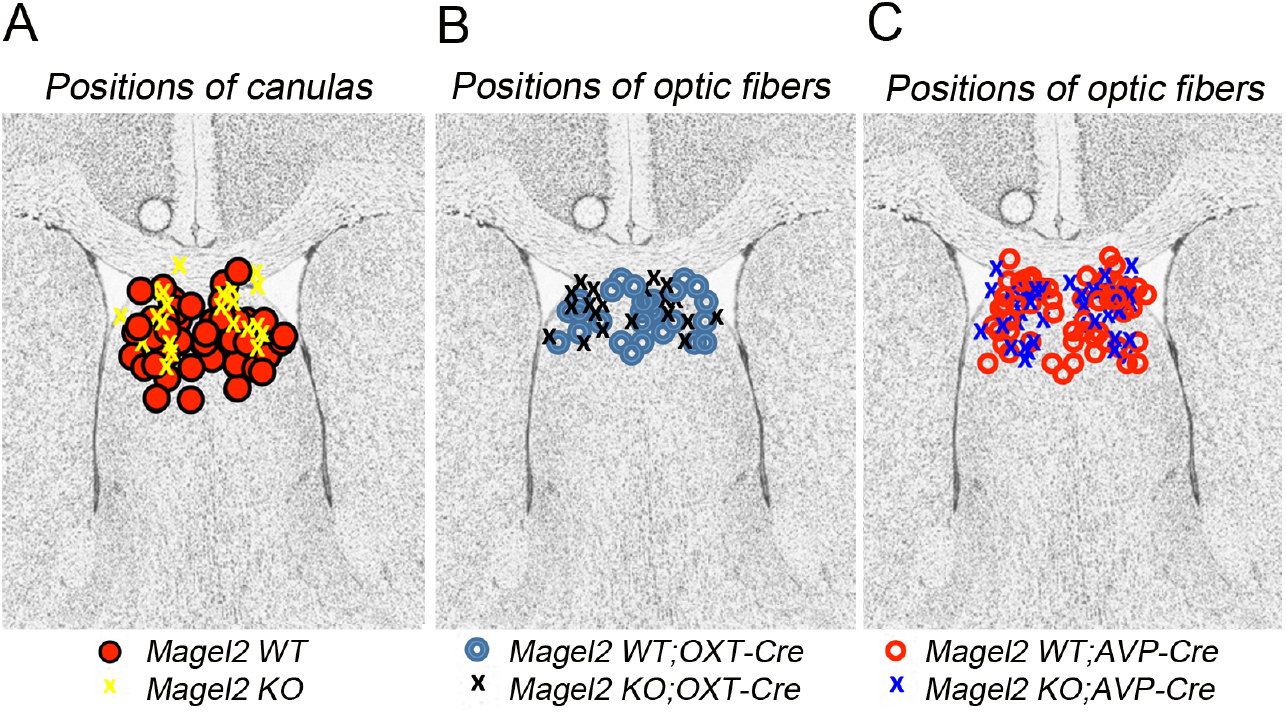
Positions of cannulas and fibers optics in septum of mice included in the study. (**a**) Postmortem histological control of the tip of cannulas on each side of LS. (**b**) Postmortem histological control of the tip of fiber optics on each side of LS in *Oxt*-*CRE* mice. (**c**) Postmortem histological control of the tip of fiber optics on each side of L

